# Post-Weaning Gut Microbiota Colonization Reveals Divergent Recovery of Skeletal Muscle and Peripheral Nerves

**DOI:** 10.64898/2026.09.07.747293

**Authors:** Sonia Calabrò, Davide Pellegrino, Chiara Cicconetti, Svenja Kankowski, Sajjad Farzin, Francesca Bertone, Mira Frühwein, Francesca Anselmi, Dario Bizzotto, Marijana Basic, Silvia Bolsega, Matthias Steglich, Lutz Wiehlmann, Cecilia Gelfi, Daniele Capitanio, Karin Kleigrewe, Salvatore Oliviero, Giovanna Gambarotta, Kirsten Haastert-Talini, Matilde Cescon, Giulia Ronchi

**Affiliations:** Department of Molecular Medicine, University of Padova, 35121, Padova, Italy; Department of Clinical and Biological Sciences & Neuroscience Institute Cavalieri Ottolenghi (NICO), University of Torino, Orbassano, 10043, (Torino), Italy; Department of Life Sciences and Systems Biology, University of Torino, 10123, Torino, Italy; Institute of Neuroanatomy and Cell Biology, Hannover Medical School, Carl-Neuberg-Str.1, 30625 Hannover, Lower-Saxony, Germany; Institute for Laboratory Animal Science and Central Animal Facility, Hannover Medical School, Carl-Neuberg-Str.1, 30625 Hannover, Lower-Saxony, Germany; Research Core Unit Genomics, Hannover Medical School, Carl-Neuberg-Str.1, 30625 Hannover, Lower-Saxony, Germany; Department of Biomedical Sciences for Health, University of Milano, 20133 Milano, Italy; Bavarian Center for Biomolecular Mass Spectrometry (BayBioMS), TUM School of Life Sciences, Technical University of Munich, 85354 Freising, Germany; Centre for Systems Neuroscience (ZSN), Hannover, 30559 Hannover, Lower-Saxony, Germany

**Keywords:** gut microbiota, germ-free mice, gut microbiota colonization, neuromuscular system

## Abstract

We previously demonstrated that the absence of a complex gut microbiota (CGM) impairs the postnatal development of peripheral nerves and motor targets in germ-free (GF) mice. In this study, we investigated whether establishing a complex gut microbiota after weaning could reverse these developmental alterations. To address this question, GF mice were colonized with a complex gut microbiota by co-housing with conventionally raised mice. Microbiota composition, peripheral nerve morphology and transcriptional profiles, skeletal muscle proteome, neuromuscular junction architecture and circulating metabolites were comprehensively analyzed and compared with those of GF, gnotobiotic OMM12 and CGM mice. Post-weaning colonization partially restored microbial diversity and resulted in a compositionally distinct microbial community with reduced alpha diversity and enrichment of *Duncaniella muris* strain B8. Despite successful microbial colonization, peripheral nerve abnormalities persisted, including axon hypermyelination, transcriptional alterations in sciatic nerves, elongated nodes of Ranvier, and dysregulated axon-glia interactions. In contrast, skeletal muscle defects were largely rescued, with restoration of muscle mass, normalization of proteomic profiles, recovery of metabolic and structural pathways, and reduced fragmentation of the postsynaptic neuromuscular junction, although presynaptic abnormalities persisted. These findings demonstrate that microbiota-dependent developmental alterations differ markedly in their reversibility across the neuromuscular system. Specifically, post-weaning colonization with a complex gut microbiota resulted in broad recovery of skeletal muscle but failed to rescue peripheral nerve abnormalities. Our findings provide a framework for future studies investigating how the timing of microbial colonization, microbiota composition, and microbiota-derived signals influence the reversibility of microbiota-dependent neuromuscular alterations.

**Significance Statement:** The gut microbiota plays an essential role in neuromuscular development. However, it is largely unknown whether developmental alterations caused by its absence can be reversed. Using germ-free mice colonized with a complex microbiota after weaning, we demonstrate that the extent of recovery differs markedly across the neuromuscular system. Although skeletal muscle largely regains its structural and molecular features, abnormalities in the peripheral nerves persist despite successful microbial colonization.

These findings show that microbiota-dependent developmental alterations are not equally reversible, pointing to the timing of microbial exposure and the availability of microbiota-derived signals as important variables. Our study provides a framework for identifying therapeutic windows and microbiota-derived signals that regulate neuromuscular development.

## Introduction

Maintaining a healthy (e.g. well-balanced) gut microbiota (GM) is essential for overall human health, as it influences metabolic, immune and neurological functions (1). Through the gut-organ axes, the GM communicates far beyond the intestine and contributes to the physiology of multiple organs and systems (2).

The neuromuscular system, which integrates peripheral nerves, skeletal muscle and the neuromuscular junction to control movement, can be affected by numerous inherited and acquired disorders, which often result in debilitating motor impairment (3). Because many of these conditions still lack effective treatments, identifying novel mechanisms that regulate neuromuscular development and maintenance is of considerable interest.

Recent research has highlighted the contribution of the GM to a wide range of diseases, including cancer, diabetes, cardiovascular disease and neurological disorders (4). Because the composition of the GM is unique to each individual and shaped by genetics, environment, and lifestyle (5), it also represents an attractive therapeutic target. Therefore, a better understanding of how individual microbial species and their bioactive metabolites influence host physiology could open new opportunities for microbiota-based interventions (6).

There is an increasing body of evidence supporting a close relationship between the GM and the neuromuscular system. Several studies have demonstrated that the GM influences skeletal muscle function and health (7–9), while growing evidence indicates that peripheral nerves are also regulated by microbial signals. In our previous work, we demonstrated that the absence of a complex microbiota can have a significant impact on peripheral nerves, skeletal muscle and neuromuscular junction. Furthermore, we showed that these abnormalities were not resolved by colonization with the simplified Oligo-Mouse-Microbiota (OMM12) consortium (7), leading us to wonder whether the successful establishment of a complex microbiota in previously germ-free mice could restore normal neuromuscular development in germ-free mice.

To address this question, germ-free mice were co-housed with conventionally raised animals after weaning and peripheral nerves, skeletal muscles and neuromuscular junctions were analysed in young adult animals. We found that post-weaning gut microbiota restoration resulted in markedly different outcomes across the neuromuscular system. While skeletal muscle showed extensive structural and molecular recovery, peripheral nerve abnormalities persisted despite successful microbial colonization. These findings demonstrate that microbiota-dependent developmental alterations are not equally reversible, and highlight the importance of the timing of microbiota restoration in influencing neuromuscular development.

## Results

### Post-weaning colonization establishes a distinct but successfully constituted gut microbiota

To determine whether co-housing with conventionally colonized mice, bearing a complex gut microbiota (CGM), was sufficient to introduce a complex gut microbiota in previously germ-free (GF) animals, we first characterized the fecal microbiota profile of EX-GF and CGM mice by shotgun metagenomic sequencing.

Post-weaning colonization successfully established a complex gut microbiota in EX-GF mice. Although the overall microbial composition broadly resembled that of CGM animals, the newly constituted microbiota in EX-GF mice remained compositionally distinct. This was evident in the relative abundance of various bacterial taxa, including the enrichment of the *Duncaniella muris* strain B8 (Fig. 1A).

**Figure 1.**
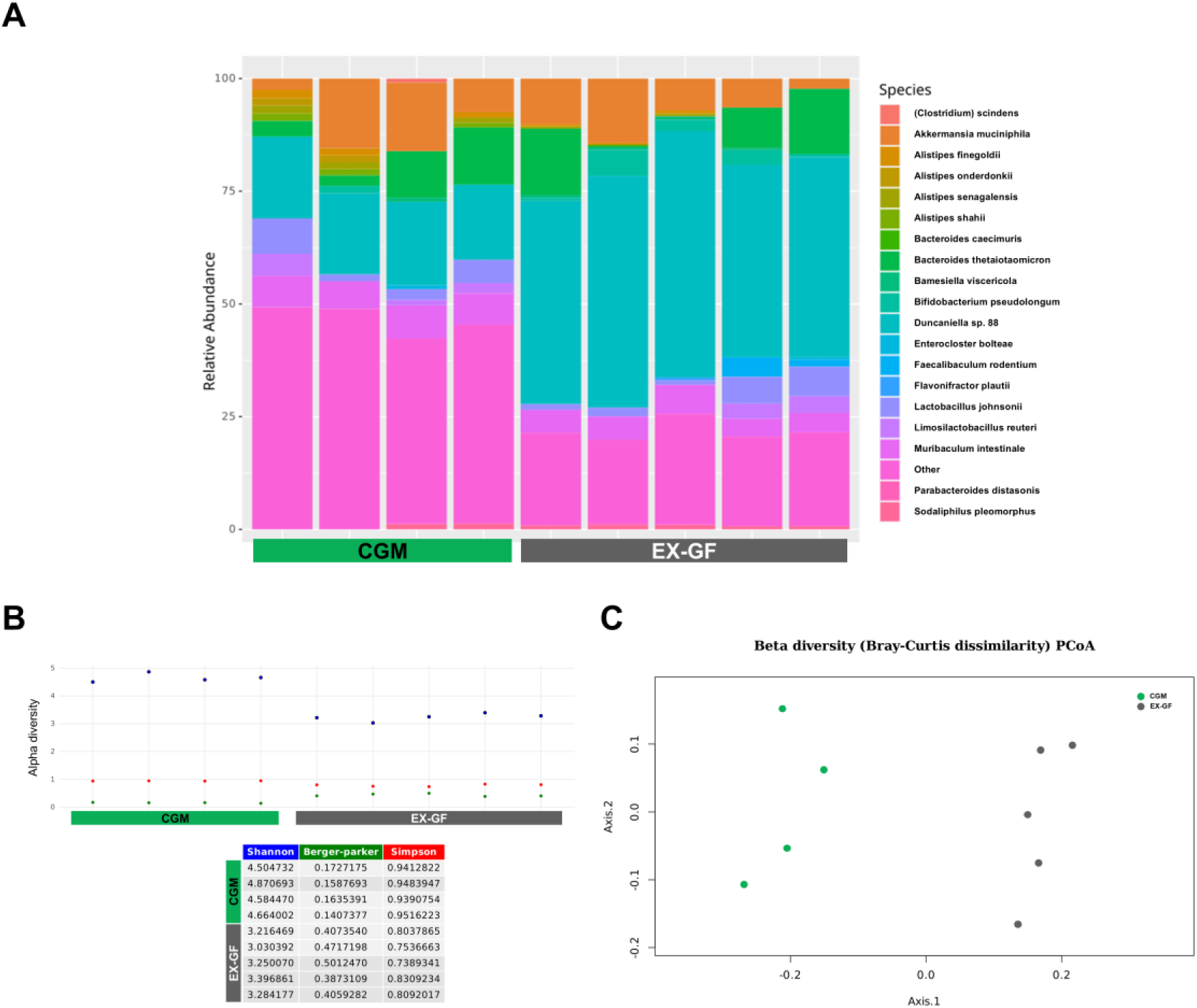
Species-level taxonomic profiling of gut microbiota in CGM and EX-GF mice. (A) Relative abundance of bacterial species identified from fecal samples of individual CGM and EX-GF mice. Taxonomic profiling was performed using shotgun metagenomics, and results are displayed as stacked bar plots representing the proportion of each species within the total microbial community per animal. (B) Alpha diversity assessed using Shannon, Berger–Parker, and Simpson indices. Each dot represents an individual sample. (C) Beta diversity analysis based on Bray–Curtis dissimilarity. Each dot represents an individual sample.

Despite successful colonization, microbial diversity was only partially transferred. Compared with CGM mice, EX-GF animals displayed significantly lower alpha diversity, reflecting reduced species richness and evenness together with greater dominance of individual taxa (Fig. 1B). Principal component analysis showed a clear separation between the microbial communities of EX-GF and CGM mice, indicating persistent differences in overall community composition following post-weaning colonization (Fig. 1C).

Taken together, these findings confirm successful post-weaning microbiota colonization and establish the experimental context for assessing the reversibility of microbiota-dependent neuromuscular alterations.

### Peripheral nerve morphology is not restored following post-weaning microbiota colonization

Next, we asked whether introducing a complex GM after weaning would be sufficient to reverse the previously identified peripheral nerve abnormalities in GF mice (7). To this end, we performed a comprehensive morphometric analysis of median nerves.

Despite successful microbiota colonization, peripheral nerve morphology was not restored in EX-GF mice. Overall, nerve architecture remained largely unchanged, with no significant differences in nerve cross-sectional area, total number of myelinated fibers, or fiber density among the experimental groups (Fig. 2A-D).

**Figure 2.**
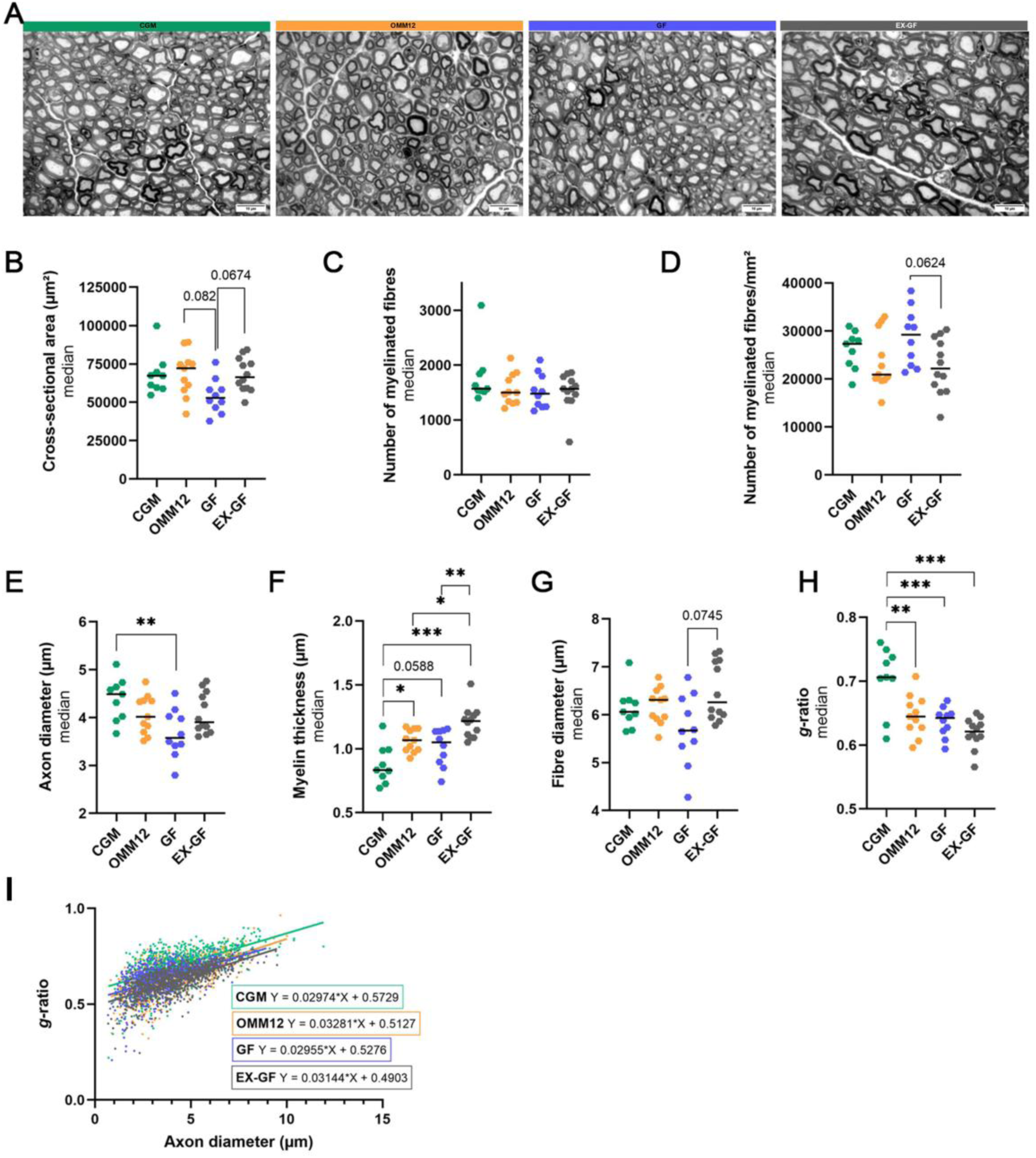
Morphoquantitative analysis of peripheral nerves. (A) Representative photomicrographs of toluidine blue stained semi-thin cross-sections. Scale bar: 10 µm. (B-D) Stereological parameters: cross-sectional area (B), total number of myelinated fibers (C), nerve fiber density (D); (E-H) morphometrical parameters: axon diameter (E), myelin thickness (F), fiber diameter (G), and g-ratio (H). Group sizes: CGM (n=9), OMM12 (n=11), GF (n=10), EX-GF (n=12). (I) Scatter plot showing g-ratio as a function of axon diameter for individual myelinated axons. Linear regression equations are provided. For (I), 560 axons were analyzed per group. Parametric data were analyzed using One-Way ANOVA followed by Tukey’s multiple comparisons post-hoc test (effect size for trend-level difference in (D) GF vs EX-GF p=0.0624 is η2 = 0.171 indicating large effect). Non-parametric data were analyzed using Kruskal-Wallis test followed by Dunn’s multiple comparisons post-hoc test (effect size for trend-level difference in (B) GF vs OMM12 p=0.0820 is r=0.381; GF vs EX-GF p=0.0674 is r = 0.391; in (G) GF vs EX-GF p=0.0745 is r = 0.386, all indicating moderate effect). * p ≤ 0.05, ** p ≤ 0.01, *** p ≤ 0.001.

In contrast, axonal and myelin morphometry revealed persistent abnormalities. Consistent with our previous findings (7), GF mice exhibited significantly smaller axon diameters than CGM mice (Fig. 2E). Myelin thickness, which was already increased in OMM12 mice, became even more pronounced following post-weaning colonization. EX-GF animals displayed the thickest myelin sheaths of all the groups (Fig. 2F). Because fiber diameter remained largely unchanged (Fig. 2G), EX-GF nerves exhibited the lowest g-ratios (Fig. 2H), indicating persistent and even exacerbated hypermyelination after delayed microbiota colonization. Scatter plot analysis further showed thicker myelin across comparable axon diameters in EX-GF nerves (Fig. 2I).

Overall, these findings demonstrate that introducing a complex gut microbiota after weaning is insufficient to normalize peripheral nerve morphology, and is instead associated with persistent and, in some parameters, exacerbated hypermyelination.

### Peripheral nerve transcriptional programs remain altered after post-weaning microbiota colonization

We next examined whether and how the transcriptional profile of peripheral nerves reflected the persistent morphological abnormalities observed after post-weaning microbiota colonization. With this aim, bulk RNA sequencing data of sciatic nerves from EX-GF mice were produced and compared to those from CGM, GF and OMM12 nerves (7).

Consistent with the morphological findings, post-weaning microbiota colonization did not restore the nerve transcriptome. Compared with CGM nerves, EX-GF nerves still displayed 339 differentially expressed genes (256 of which were upregulated and 83 downregulated; see Fig. 3A), indicating persistent transcriptional alterations despite successful microbial colonization.

**Figure 3.**
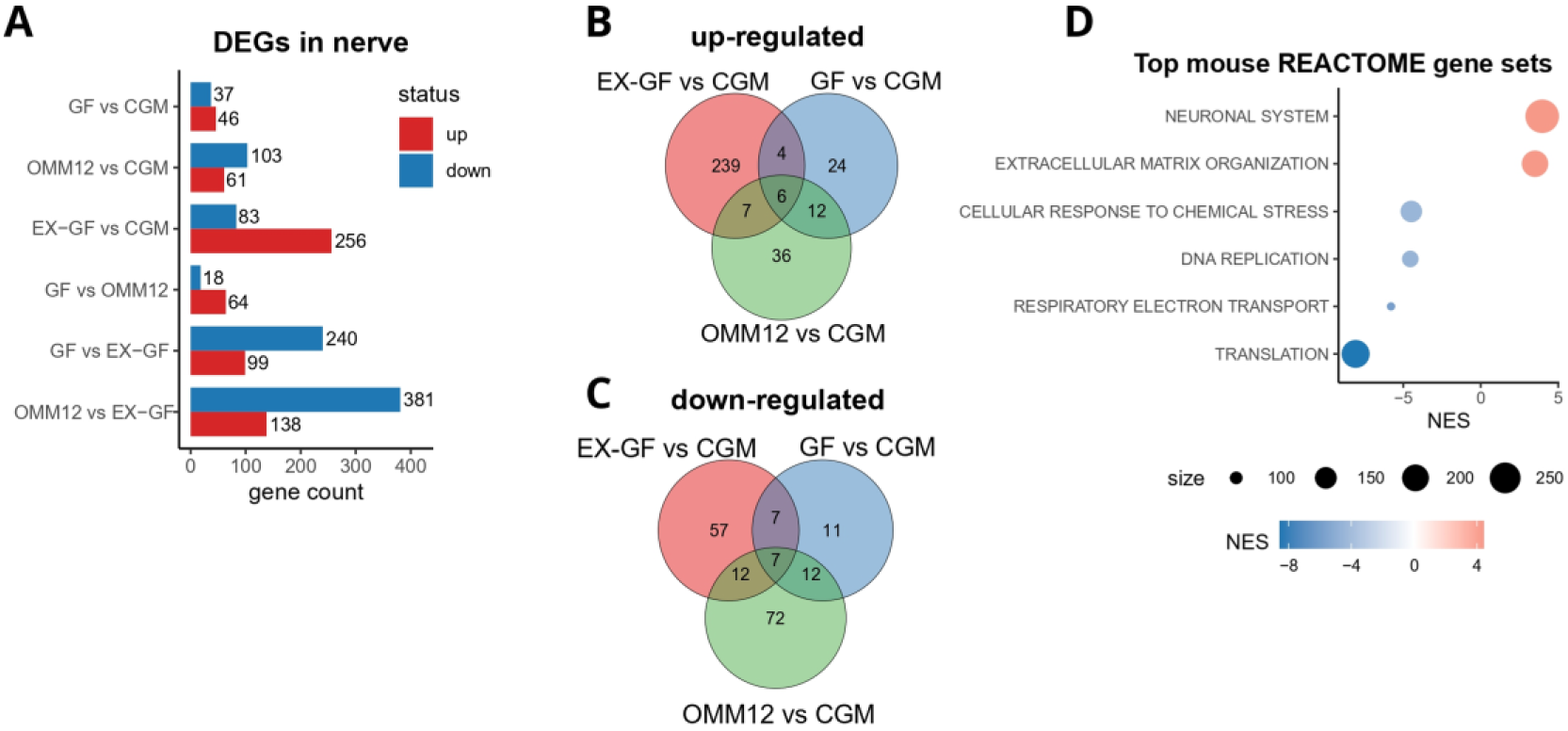
Transcriptomic profiles of sciatic nerves. (A) Count of differentially expressed genes (DEGs) across all pairwise comparisons in sciatic nerve samples. Up-regulated genes are shown in red, downregulated genes in blue. DEGs were defined by thresholds of |logFC| > 1 and p-value < 0.01. (B-C) Venn diagrams showing overlap of DEGs among comparisons in sciatic nerves. (D) Significantly enriched REACTOME pathways from GSEA. Genes were ranked according to logFC(log10p-value) comparing EX-GF vs CGM in sciatic nerve samples. Size of the dots represents the dimension of the geneset. Positive NES in red, negative in blue.

Of note, comparison of differentially expressed genes across experimental groups further revealed that many transcriptional changes observed in EX-GF nerves were unique to this condition, with only limited overlap with GF or OMM12 mice (Fig. 3B, C). These findings suggest that delayed, e.g. post-weaning, microbiota colonization accompanies with a distinct transcriptional profile rather than the physiological one.

Gene set enrichment analysis (GSEA) identified positive enrichment of REACTOME pathways related to neuronal system function and extracellular matrix organization, whereas pathways involved in cellular responses to chemical stress, DNA replication, respiratory electron transport and translation were negatively enriched (Fig. 3D), indicating persistent remodelling of neuronal and metabolic programs following post-weaning microbiota colonization.

Together, these findings demonstrate that restoration of a complex gut microbiota after weaning is insufficient to normalize peripheral nerve transcriptional programs, consistent with the persistent structural abnormalities observed in EX-GF nerves.

### Persistent structural abnormalities are accompanied by altered axon-glia molecular programs

To investigate the structural and molecular basis of the persistent hypermyelination observed in EX-GF nerves, we next examined myelin ultrastructure together with the expression of key regulators of peripheral myelination and nodal organization.

Electron microscopy showed preserved myelin compaction across all experimental groups, indicating that the ultrastructural organization of compact myelin remained intact (Fig. 4A). By contrast, GF, OMM12 and EX-GF nerves displayed a significantly higher number of myelin lamellae than CGM nerves (Fig. 4B), indicating that hypermyelination results from excessive Schwann cell wrapping rather than altered myelin compaction (Fig. 4C).

**Figure 4.**
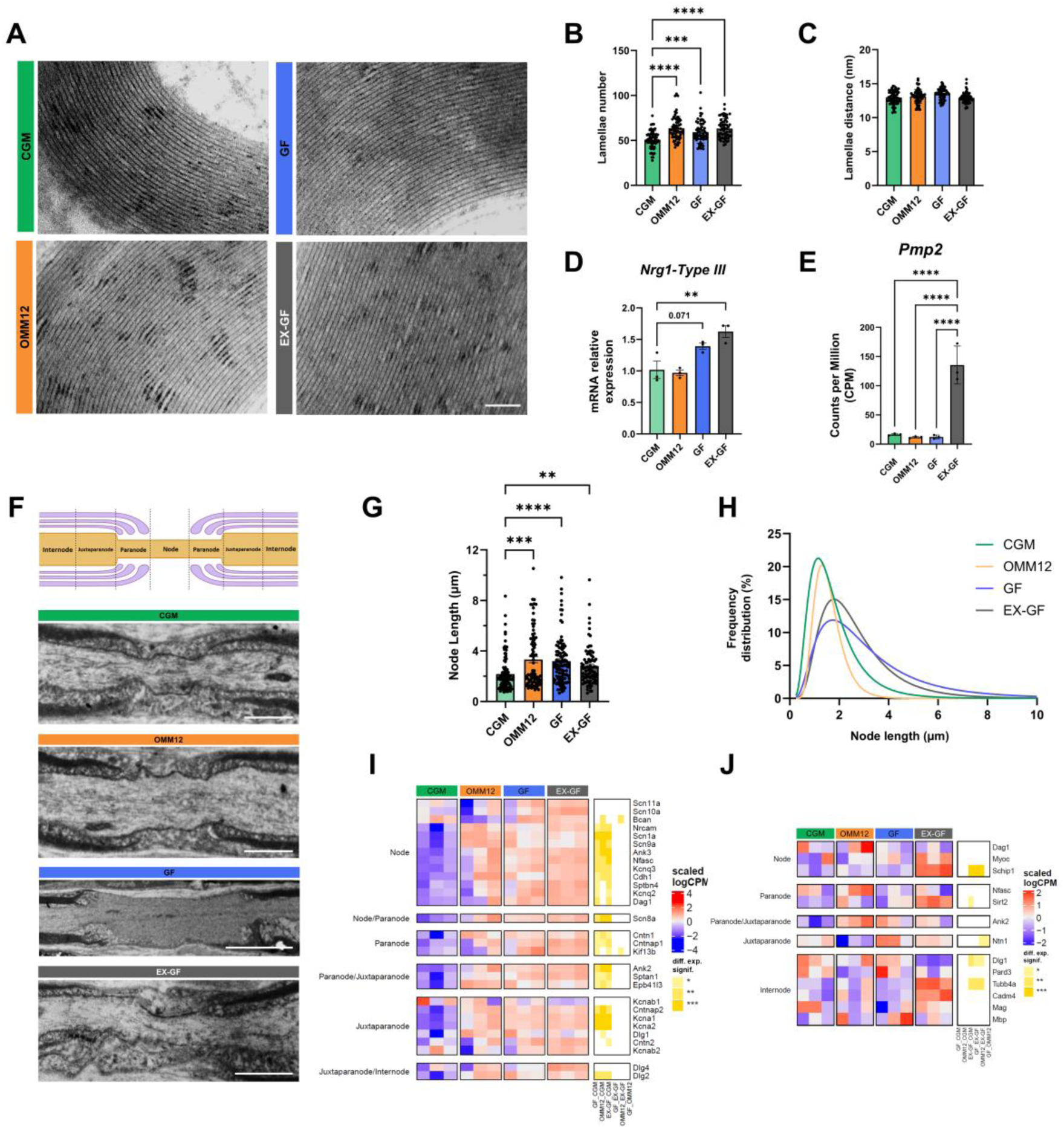
Analysis of myelinated nerve fibers. (A) Representative electron microscopy images of ultrathin transverse sections showing myelin lamellae in median nerves. Scale bar = 0.1µm; (B, C) Quantification of myelin structure by EM: (B) Number of lamellae, (C) Interlamellar spacing (lamellae distance). Data represent mean ± SEM from 60 fibers per group. One-way ANOVA; *p ≤ 0.001, **p ≤ 0.0001, ***p ≤ 0.00001, ****p ≤ 0.000001. (D) qRT-PCR analysis of *Nrg1-Type III* transcript levels. One-way ANOVA; **p ≤ 0.01 (n=3 per group). (E) Gene expression of *Pmp2* shown as counts per million (CPM). One-way ANOVA; *p ≤ 0.05, **p ≤ 0.01, ***p ≤ 0.001, ****p ≤ 0.0001 (n = 3 per group). (F) Schematic of nodal domain regions followed by representative EM images of nodes of Ranvier in median nerves from each group. Scale bar CGM, OMM12 and EX- GF: 1µm; GF: 5 µm; (G) Quantification of nodal length from EM images. Data represent mean± SEM. One-way ANOVA; *p ≤ 0.05, **p ≤ 0.01, ***p ≤ 0.001, ****p ≤ 0.0001 (CGM=109 nodes, OMM12=80, GF=105, EX-GF=86). (H) Graphical representation of node of Ranvier length. Curves were fitted to data using non-linear regression (Lognormal). (I-J) Heatmaps of expressed genes associated with nodal domains (node, node/paranode, paranode/juxtaparanode, juxtaparanode, or juxtaparanode/internode) in DRG samples (I) and sciatic nerve samples (J). Scaled log-expression values from 3 samples of each experimental group. Red and blue indicate high and low expression, respectively. Differential expression significance indicates statistical significance in each pairwise comparison (*p ≤ 0.05, **p ≤ 0.01, ***p ≤ 0.001).

To investigate the molecular basis of persistent hypermyelination, we measured the expression of Neuregulin-1 type III, a major regulator of peripheral myelination (10, 11), in dorsal root ganglia by real-time qPCR. *Nrg1* was significantly upregulated in EX-GF mice and showed a similar trend in GF mice (Fig. 4D). Moreover, we analyzed RNA-seq data focusing on the “myelin sheath” GO term (GO:0043209). Among the few identified DEGs (Supplementary Fig. S1A), Peripheral myelin protein 2 (*Pmp2*) was significantly upregulated in EX-GF compared to CGM (Fig. 4E). Aberrant Schwann cell-axon interactions, including myelin outfoldings, myelinated inclusions and myelin splitting, were also observed in OMM12, GF and EX-GF nerves (Supplementary Fig. S1B).

Persistent hypermyelination was not limited to compact myelin but was accompanied by additional alterations of myelinated nerve fibers. While internodal length remained largely unchanged across experimental groups (Supplementary Fig. S1C, D), nodes of Ranvier were significantly longer in OMM12, GF and EX-GF mice than in CGM controls (Fig. 4F- H). These structural alterations were paralleled by a widespread dysregulation of node-associated genes, predominantly involving axonal components, whereas relatively few Schwann cell-associated genes were altered (Fig. 4I,J).

Together, these observations indicate that delayed microbiota colonization fails to restore the molecular programs governing axon-Schwann cell interactions.

### Skeletal muscle development is largely restored following post-weaning microbiota colonization

We next asked whether skeletal muscle responded differently than peripheral nerves to post-weaning microbiota colonization. Gastrocnemius (GA) muscle weight, significantly reduced in GF mice, returned to CGM levels in EX-GF animals (Fig. 5A). Similar results were observed in tibialis anterior (TA) muscle (Supplementary Fig. S3A).

**Figure 5.**
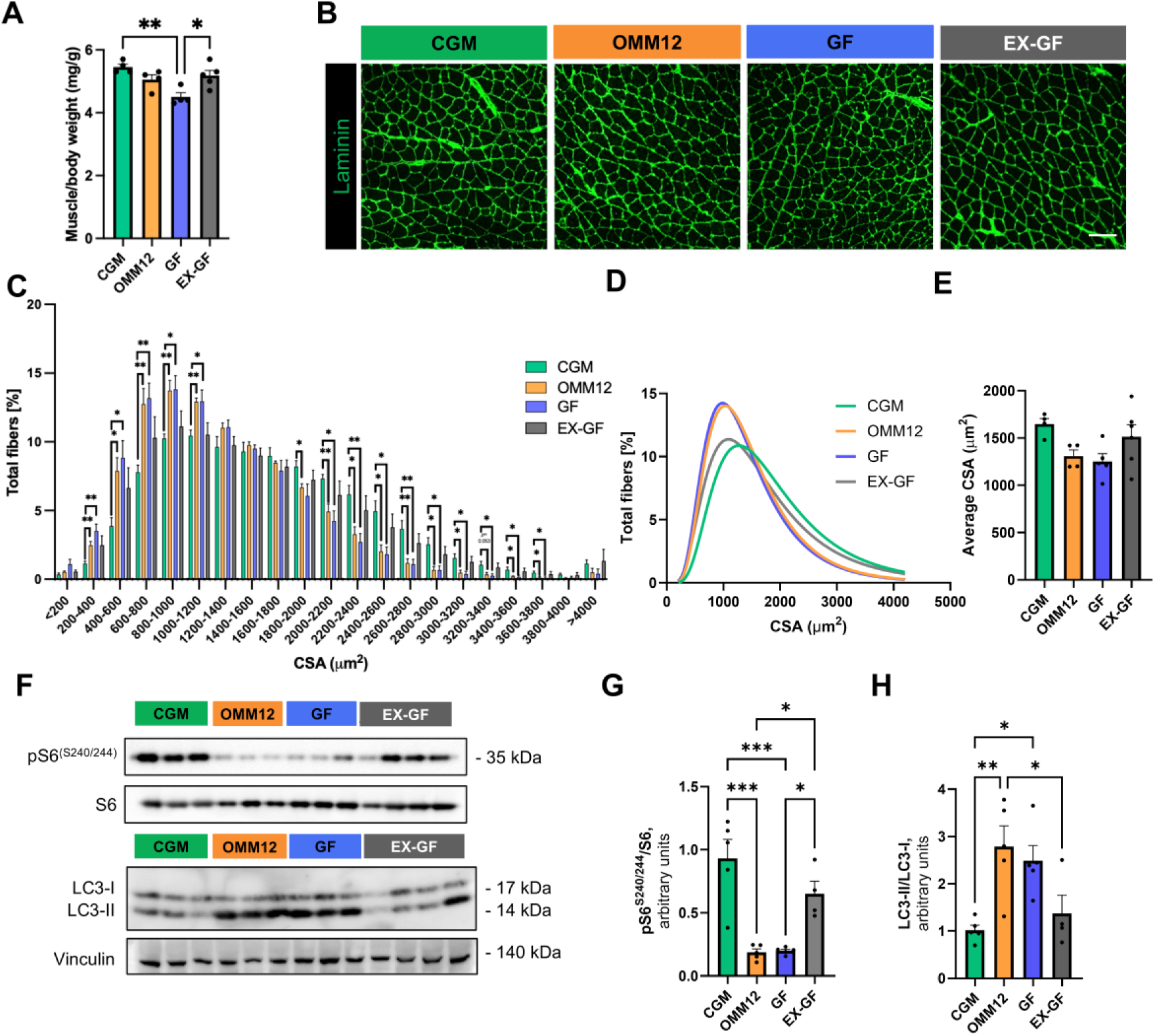
Morphometric and molecular analysis of skeletal muscle. (A) Gastrocnemius (GA) muscle weight normalized by mice body weight in the different groups (*p ≤ 0.05; **p ≤ 0.01; one-way ANOVA with post hoc Tukey’s multiple comparisons test; CGM, OMM12, GF, n=4; EX-GF, n=5). (B) Representative fluorescence micrographs of GA cross-sections from CGM, OMM12, GF and EX-GF mice, stained with laminin (green), marking fiber edges. Scale bar: 100 µm. (C) Comparison of myofiber CSA distribution among CGM, OMM12, GF and EX-GF GA muscles (*p ≤ 0.05; **p ≤ 0.01; multiple unpaired two-tailed Student’s t-tests; CGM, OMM12, n=4; GF, n=5; EX-GF, n=6). (D) Graphical representation of myofiber CSA distribution in GA muscles from the different groups. Curves were fitted to data using non-linear regression (Lognormal). (E) Quantification of average myofiber cross-sectional area (CSA) in GA muscles from CGM, OMM12, GF and EX-GF mice (one-way ANOVA with Tukey’s post hoc test for multiple comparisons; CGM, OMM12, n=4; GF, n=5; EX-GF, n=6). (F) Representative images of western blot analysis of LC3 lipidation and S6 phosphorylation in total protein lysates of GA muscles from CGM, OMM12, GF and EX-GF mice. (G, H) Densitometric quantifications of pS6 normalized to S6 (G) and of LC3-II normalized to LC3-I (H) (*p ≤ 0.05; **p ≤ 0.01; ***p ≤ 0.001; one-way ANOVA with post hoc Tukey’s multiple comparisons test; CGM, OMM12, GF n=5; EX-GF, n=4).

Morphometric analysis of GA cross-sections (Fig. 5B) showed that GF muscles displayed a clear left shift in both cross-sectional area (CSA) and minimum Feret’s diameter (MFD) distributions, indicating an increased proportion of smaller myofibers (Fig. 5C-E; Supplementary Fig. S2A-C). In contrast, the CSA and MFD distributions of EX-GF muscles almost completely overlapped with those of CGM mice, demonstrating morphological recovery of myofiber size (Fig. 5C-E; Supplementary Fig. S2A-C). Interestingly, OMM12 mice exhibited a markedly different phenotype. Although muscle weight was comparable to CGM animals (Fig. 5A), their CSA and MFD distributions remained similar to those of GF mice and significantly differed from CGM controls (Fig. 5C-E; Supplementary Fig. S2A-C).

A similar, although less remarkable, pattern was observed in the TA cross-sections, where reductions in myofiber size detected in GF mice were largely restored following post-weaning microbiota colonization (Supplementary Fig. S3). However, differently from what was observed in GA, OMM12 colonization was also sufficient to normalize TA morphometric parameters (Supplementary Fig. S3), suggesting distinct responses of individual skeletal muscles to simplified microbial communities.

Together, these findings demonstrate that restoring a complex gut microbiota after weaning is sufficient to reverse skeletal muscle atrophy and morphological abnormalities, suggesting a causative role for GM in the regulation of skeletal muscle mass.

### Skeletal muscle protein homeostasis is restored following post-weaning microbiota colonization

To investigate the molecular bases underlying the recovery of skeletal muscle morphology, we next examined key signaling pathways regulating protein turnover, as skeletal muscle mass is determined by the balance between protein synthesis and protein degradation (7, 8).Consistently with morphological results, expression of the atrogenes *Tripartite Motif Containing 63* (*Trim63*, widely known as *MuRF1*) and *F-Box Protein 32* (*Fbxo32*, or *Atrogin-1*) was increased in GF and OMM12 GA muscles and returned to control levels following post-weaning microbiota colonization (Supplementary Fig. S2D, E).

Moreover, reduced phosphorylation of ribosomal protein S6 (on S240/244), a downstream target of Akt/mTOR signaling controlling protein synthesis, and increased LC3-II levels, reflecting enhanced autophagic activity, were both normalized following post-weaning microbiota colonization of GF mice (Fig. 5F-H). A comparable rescue was also observed in TA muscles (Supplementary Fig. S4A-C).

Together, these findings suggest that introducing a complex gut microbiota in previously GF mice is sufficient to normalize the major molecular pathways regulating skeletal muscle protein homeostasis in EX-GF mice.

### Post-weaning microbiota colonization broadly normalizes the skeletal muscle proteome

We next asked whether normalization of protein turnover was accompanied by remodeling of the skeletal muscle proteome. To address this, we performed unbiased label-free LC-MS/MS proteomic analysis of GA muscles.

The analysis identified widespread alterations in protein abundance in both GF and OMM12 muscles, whereas the EX-GF proteome closely resembled that of CGM mice.

Compared with CGM, 263 and 276 proteins were differentially expressed in GF and OMM12 muscles, respectively, whereas only 63 proteins differed in EX-GF muscles (Fig. 6A). Consistent with these observations, hierarchical clustering grouped EX-GF samples together with CGM, clearly separating them from GF and OMM12 muscles (Fig. 6B).

**Figure 6.**
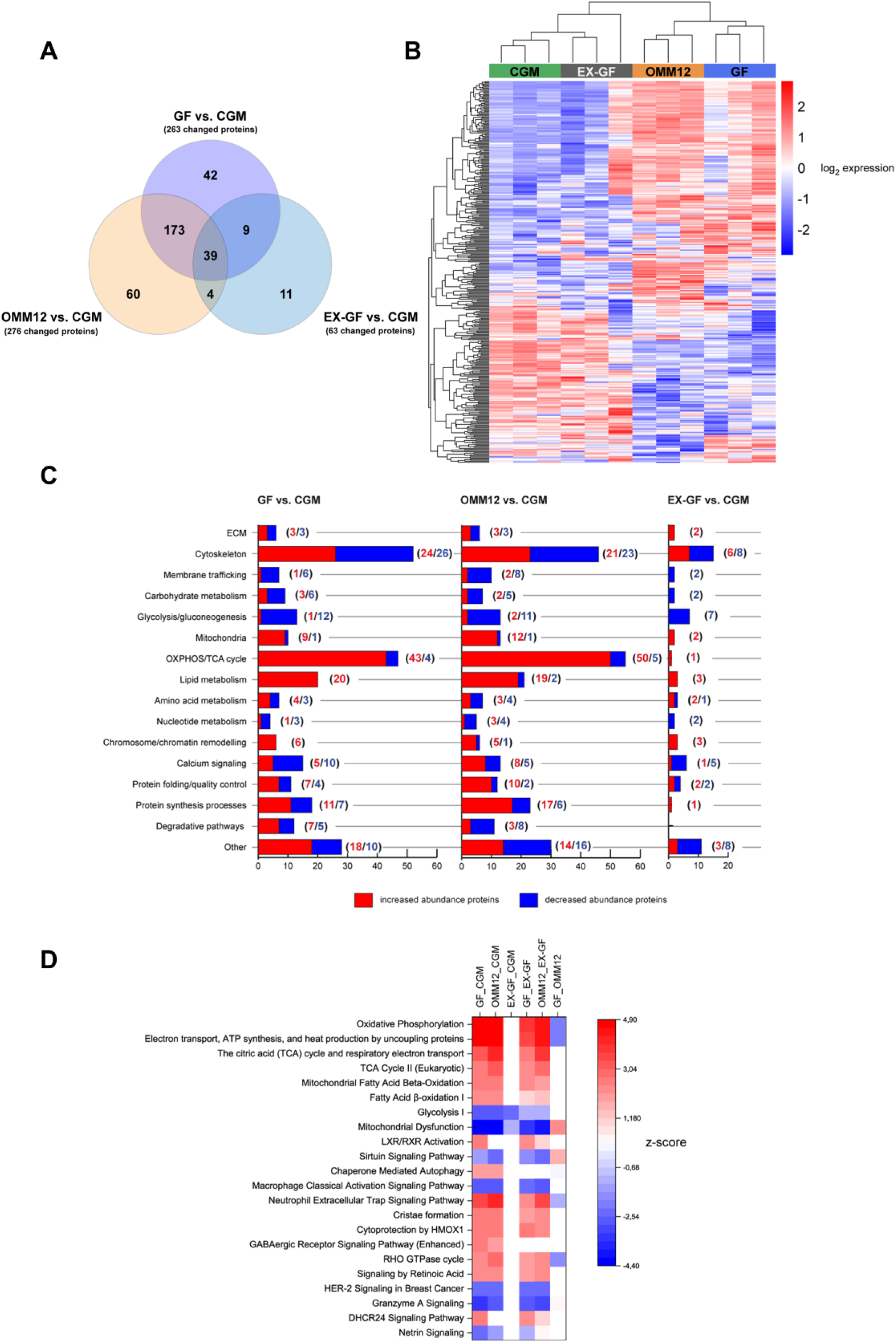
Proteomic profiles of gastrocnemius muscles. (A) Venn diagram comparing the number of differentially expressed proteins in gastrocnemius (GA) muscles from GF, OMM12 and EX-GF mice compared to CGM controls. (B) Comparison of the proteome profiles from CGM, OMM12, GF and EX-GF GA muscles analysed by unsupervised hierarchical clustering performed using Euclidean distance matrix and average linkage method. (C) Functional annotation of upregulated and downregulated proteins expressed in GA muscles from GF, OMM12 and EX-GF mice compared to CGM controls. (D) Heatmap of canonical pathways predicted to be significantly enriched (red) or inhibited (blue) based on protein changes in GF, OMM12 and EX-GF muscles, as determined by Ingenuity Pathway Analysis (a Fisher’s exact test p-value < 0.05 and z-scores ≤ −2 and ≥ 2 were considered as statistically significant difference; n=3 mice, each group).

Functional classification of dysregulated proteins revealed that the largest changes in GF and OMM12 muscles involved proteins associated with oxidative phosphorylation / tricarboxylic acid (TCA) cycle and cytoskeletal organization. These alterations were largely absent in EX-GF muscles (Fig. 6C). Proteins involved in protein synthesis and degradative pathways were also markedly altered in GF and OMM12 muscles but were largely normalized following post-weaning microbiota colonization (Fig. 6C).

Ingenuity Pathway Analysis further supported these observations. Compared to CGM, GF and OMM12 muscles displayed largely shared deregulated pathway signatures, characterized by activation of oxidative phosphorylation, respiratory electron transport, the TCA cycle and fatty acid β-oxidation, together with inhibition of glycolysis and mitochondrial dysfunction, suggesting a shift toward pro-oxidative metabolism (Fig. 6D). With the only exception of glycolysis, which remained inactivated, all these pathways were not altered in EX-GF muscles, whose proteomic profile closely resembled that of CGM animals, (Fig. 6D).

Collectively, these findings demonstrate that establishing of a complex GM after weaning is accompanied by a near-complete normalization of the aberrant GF muscle proteome, suggesting a direct modulation of GM of skeletal muscle protein composition

### Proteome normalization is accompanied by recovery of skeletal muscle metabolism

Proteomic analysis identified oxidative phosphorylation and TCA cycle proteins as the most extensively deregulated metabolic category in GF and OMM12 muscles (Fig. 7A). Nearly all of these alterations were absent in EX-GF muscles.

**Figure 7.**
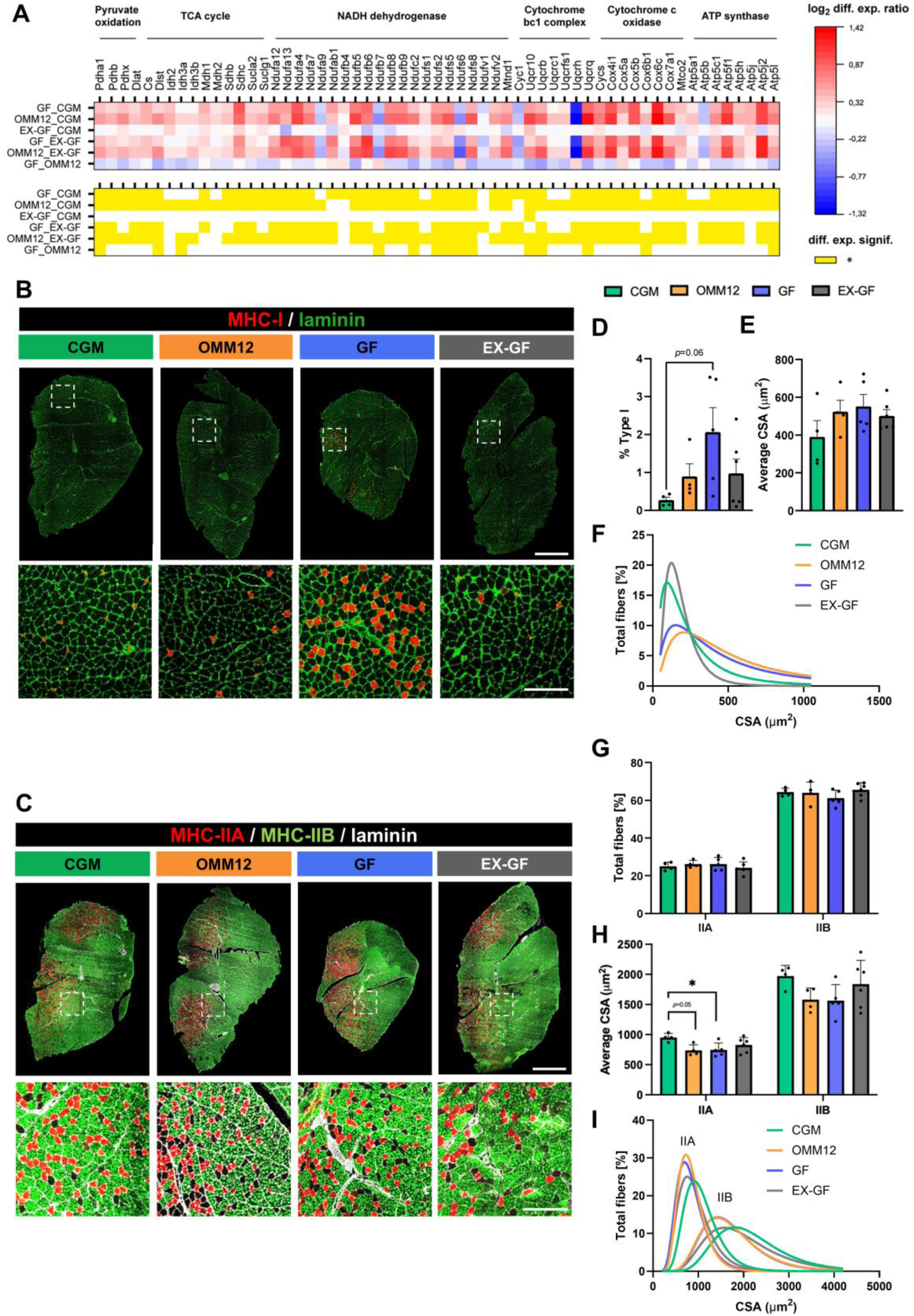
Analysis of metabolic changes in gastrocnemius muscles upon microbiota modulation. (A) Heatmaps of differentially expressed proteins associated with the OXPHOS/TCA cycle functional annotation. Each column represents a protein associated to specialized processes or enzymatic complexes (pyruvate oxidation, TCA cycle, NADH dehydrogenase, cytochrome bc1 complex, cytochrome c oxidase and ATP synthase); each row corresponds to the indicated pairwise comparison between experimental groups. Color intensity reflects the log2-scaled average differential expression ratio between the two groups; red = upregulated, blue = downregulated. Statistical significance is indicated in yellow in corresponding columns (*p ≤ 0.05, one-way ANOVA test with Tukey’s post hoc test for multiple comparisons and Benjamini-Hochberg FDR). (B, C) Representative fluorescence microscopy images of whole GA cross-sections from CGM, OMM12, GF and EX-GF mice, stained with antibodies against MHC-I (red) and laminin (green) (B) or with MHC-IIA (red), MHC-IIB (green) and laminin (gray) (C). Dotted white squares indicate the areas that are shown at higher magnification in the bottom panels. Scale bars: 1000 μm (top panels) or 200 μm (bottom panels). (D, E) Quantification of the percentages of type I fibers (D) and of their average cross-sectional area (CSA) in GA muscles from CGM, OMM12, GF and EX-GF mice, based on immunofluorescent images as in (B) (one-way ANOVA test with Tukey’s post hoc test for multiple comparisons; CGM, OMM12, n=4; GF, n=5; EX-GF, n=6). (F) Graphical representation of type I myofiber CSA distribution in GA muscles from the different groups. Curves were fitted to data using non-linear regression (Lognormal). (G, H) Quantification of the percentages of type IIA and IIB fibers (G) and of their respective average CSA in GA muscles from CGM, OMM12, GF and EX- GF mice, based on immunofluorescent images as in (C) (*p ≤ 0.05, one-way ANOVA test with Tukey’s post hoc test for multiple comparisons; CGM, OMM12, n=4; GF, n=5; EX-GF, n=6). (I) Graphical representation of type IIA and IIB myofiber CSA distribution in GA muscles from the different groups. Curves were fitted to data using non-linear regression (Lognormal). Data are presented as mean ± SEM.

To understand whether these molecular changes were reflected by an altered metabolic capacity at the tissue level, we investigated fiber type composition in the GA muscle, a predominantly fast-twitch mixed muscle, by immunostaining for myosin heavy chain (MHC) MHC-I, MHC-IIA and MHC-IIB fiber subtypes (Fig. 7B, C). GF, but not OMM12 muscles, showed a clearly higher proportion of oxidative type I fibers than CGM muscles, whereas this increase was no longer observed following post-weaning microbiota colonization (Fig. 7D). Although the average size of type I fibers was largely unchanged in terms of average CSA and MFD (Fig. 7E), their size distribution shifted toward larger fibers in both GF and OMM12 muscles and returned to a CGM-like profile in EX-GF animals (Fig. 7F; Supplementary Fig. S5A, B).

The overall proportion of type IIA and type IIB fibers, on the contrary, was not significantly affected by microbiota composition (Fig. 7G). However, type IIA and type IIB fibers from GF and OMM12 muscles displayed a significant or a trend to reduction in fiber size, respectively, which was no longer evident in EX-GF muscles (Fig. 7H, I; Supplementary Fig. S5C, D), suggesting that fast-twitch, more than slow-twitch fibers, are susceptible to atrophy induced by GM absence. Despite these morphological changes, expression of MHC isoforms remained unchanged across all experimental groups (Supplementary Fig. S5E).

Together, these data indicate that the metabolic remodeling induced by the absence of a complex GM is reversible following post-weaning microbiota colonization.

### Microbiota colonization restores contractile architecture of skeletal muscle

Besides the metabolic “OXPHOS/TCA cycle” class, proteomic analysis identified cytoskeletal proteins as the second largest class of dysregulated proteins in GF and OMM12 muscles (Fig. 6C). Among them, many proteins were structural components of the sarcomere, the functional contractile unit of skeletal muscle (Supplementary Fig. S6A), associated with thick and thin filaments, thin filament length regulation, the Z-disc and the M-band. Most of these proteins returned to CGM levels following post-weaning microbiota colonization, including 4 of 5 proteins involved in thin filament length regulation, 12 of 15 Z-disc proteins and all M-band-associated proteins (Fig. 8A).

**Figure 8.**
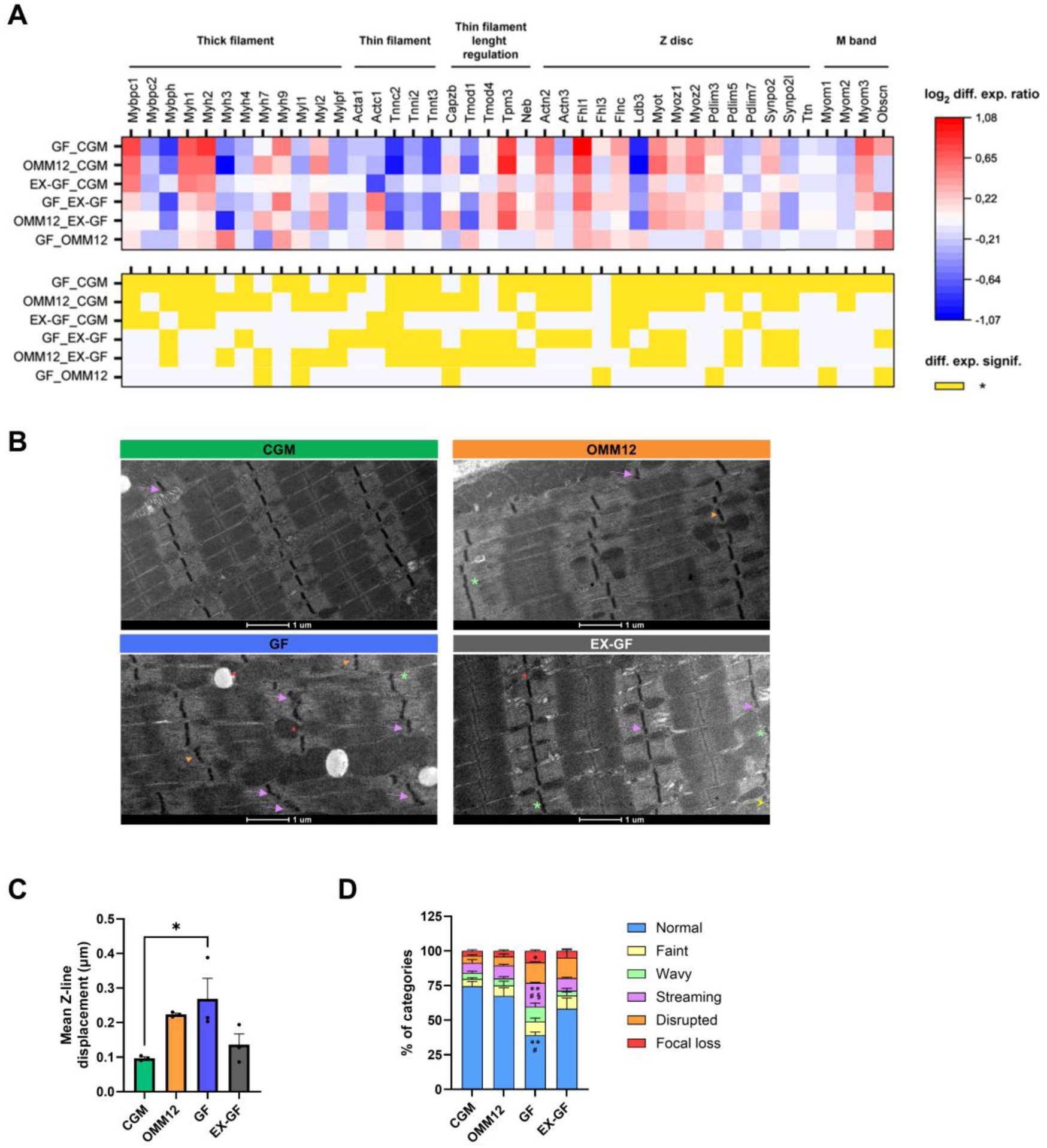
Analysis of sarcomeric alterations in diaphragm muscles upon microbiota modulation. (A) Heatmaps of differentially expressed proteins annotated in the functional class “cytoskeleton” and associated with the sarcomere. Each column represents a protein associated with specialized structures or regulatory aspects of the sarcomere (thick filaments, thin filaments, thin filament length regulation, Z-disc, M band); each row corresponds to the indicated pairwise comparison between experimental groups. Color intensity reflects the log2-scaled average differential expression ratio between the two groups; red = upregulated, blue = downregulated. Statistical significance is indicated in yellow in corresponding columns (*p ≤ 0.05, one-way ANOVA test with Tukey’s post hoc test for multiple comparisons and Benjamini-Hochberg FDR). (B) Representative electron microscopy images of longitudinal sections of diaphragms. Z-disc lines with faint (yellow dotted arrow), wavy (green asterisk), streaming (purple arrow) or disrupted (orange arrowhead) appearance as well as Z line focal loss (red star) are indicated. Scale bar: 1 μm. (C) Quantification of mean longitudinal Z-line displacement in diaphragms from CGM, OMM12, GF and EX-GF mice, analyzed through TEM images as in (B). Data represent mean ± SEM (*p ≤ 0.05, one-way ANOVA test with Tukey’s post hoc test for multiple comparisons; n=3, per group). (D) Quantification of Z-line phenotypic appearance, classified as normal, faint, wavy, streaming, disrupted and focal loss. (*p ≤ 0.05; **p ≤ 0.01; one-way ANOVA test with Tukey’s post hoc test for multiple comparisons; *: CGM vs GF; #: OMM12 vs GF; §: EX-GF vs GF; n=3, each group). Data are shown as mean ± SEM.

Despite such alterations, ultrastructural analysis of longitudinal diaphragm sections revealed preserved overall sarcomere architecture across all experimental groups, with no differences in sarcomere length, A-band, I-band or Z-line dimensions (Supplementary Fig. S6B-G). Nonetheless, GF muscles displayed misalignment of Z-lines compared to CGM muscles (Fig. 8B, C; Supplementary Fig. S7A). GF muscles were also characterized by a higher frequency of phenotypically abnormal Z-lines, including faint, wavy, streaming, disrupted and focally lost Z-discs (Fig. 8B, D; Supplementary Fig. S7B). Notably, both abnormalities were reversed in EX-GF mice.

Together, these findings indicate that developing a complex gut microbiota after weaning also normalizes the structural organization of the contractile apparatus.

### Neuromuscular junctions exhibit differential recovery of pre- and postsynaptic compartments following microbiota colonization

We next asked whether the differential recovery observed in peripheral nerves and skeletal muscle was also reflected at the neuromuscular junction (NMJ), where the two compartments functionally interact.

As previously reported (7), GF diaphragms exhibited significant fragmentation of acetylcholine receptor (AChR) clusters compared to CGM mice (see Fig. 9A–D). The number of AChR fragments per NMJ and the fragmentation index were both significantly higher in GF animals, but returned to CGM levels following post-weaning microbiota colonization in EX-GF mice. A similar condition was also observed in OMM12 mice (Fig. 9B–D), indicating that postsynaptic organization depends largely on the presence of microbiota, even if it is modified or simplified.

**Figure 9.**
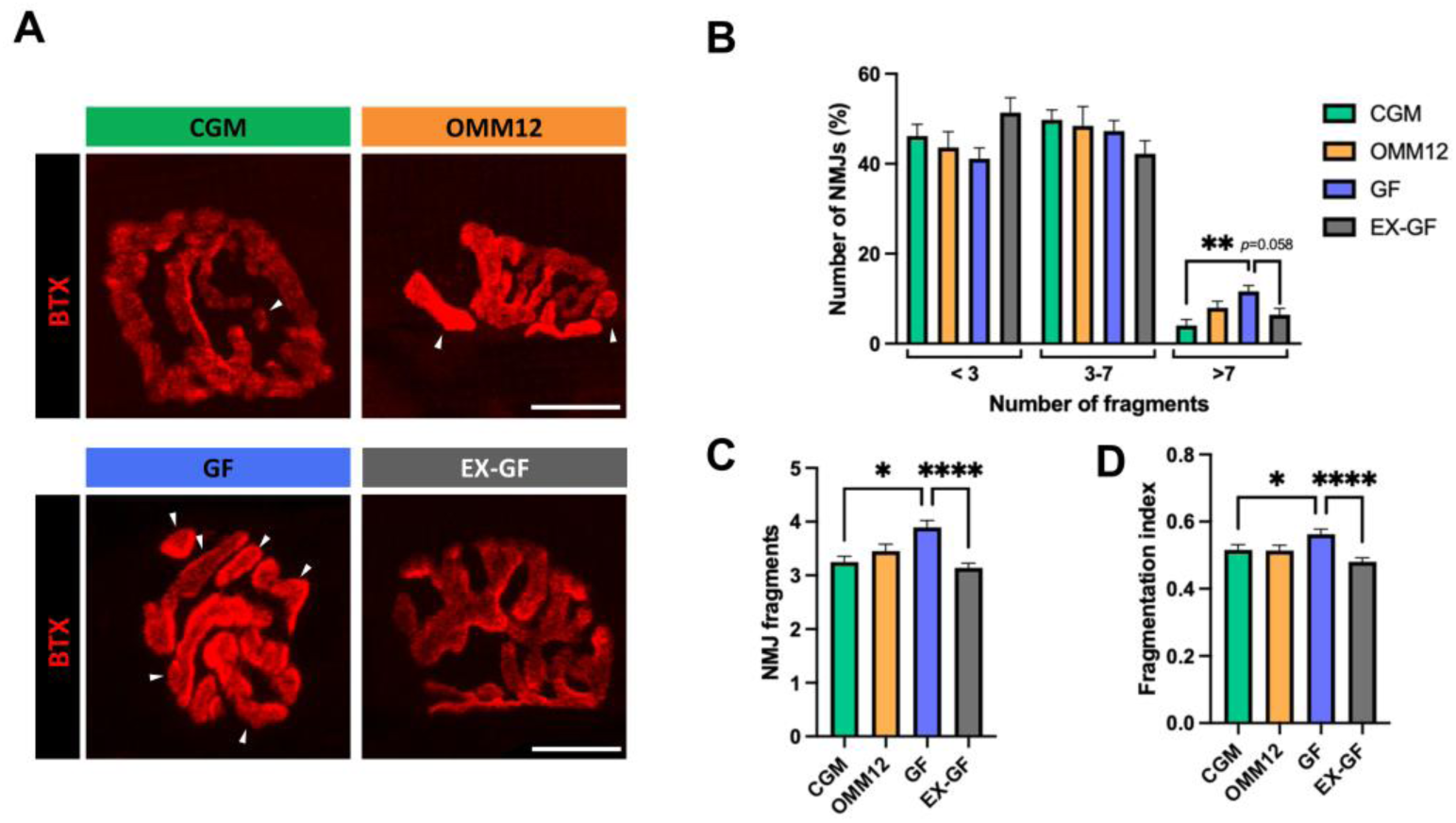
Morphometric analysis of NMJ. (A) Representative images of α-bungarotoxin-stained NMJs (red) in diaphragms. White arrowheads indicate isolated AChR clusters. Scale bar: 10 μm. (B) Percentage of total NMJs presenting the reported number of fragments (**p ≤ 0.01; one-way ANOVA test with Tukey’s post hoc test for multiple comparisons; n=8 mice, each group). (C, D) Analysis of postsynaptic fragmentation, as average number of fragments per NMJ (C) and fragmentation index (D) (*p ≤ 0.05; ****p ≤ 0.0001; Kruskal–Wallis test with post-hoc Dunn’s test for multiple comparisons; CGM n=414; OMM12 n=447; GF n=475; EX-GF n=724 NMJs, sampled from 8 mice). Data are depicted as mean ± SEM.

To further characterize NMJ architecture, whole-mount diaphragm preparations were stained for postsynaptic acetylcholine receptors together with presynaptic and axonal markers, allowing quantitative morphometric analysis of both synaptic compartments using the aNMJ-morph macro (12) (Supplementary Fig. S8A). Postsynaptic AChR and endplate morphology were largely preserved across experimental groups, although OMM12 diaphragms displayed smaller endplates than the other groups (Supplementary Fig. S8B-F). At the presynaptic side, while no differences were detected in nerve terminal perimeter or complexity across groups, some abnormalities were noted in GF NMJs, including a reduced axon diameter (Supplementary Fig. S8G), and an enlarged nerve terminal area (Supplementary Fig. S8I), that persisted in EX-GF samples. Both GF and EX-GF muscles also showed increased overlap between pre- and postsynaptic compartments (Supplementary Fig. S8K), despite recovery of postsynaptic fragmentation. These findings mirror the distinct responses of skeletal muscle and peripheral nerves to post-weaning microbiota colonization, with recovery of the postsynaptic compartment but persistence of presynaptic abnormalities.

### Distinct microbial and metabolic signatures accompany neuromuscular recovery following post-weaning microbiota colonization

Finally, we investigated whether the differential recovery observed across the neuromuscular system was accompanied by distinct systemic metabolic signatures. Targeted metabolomic analysis of serum samples revealed clearly separated metabolic profiles among the experimental groups, with EX-GF mice clustering closer to CGM animals than to GF or OMM12 mice (Fig. 10A).

**Figure 10.**
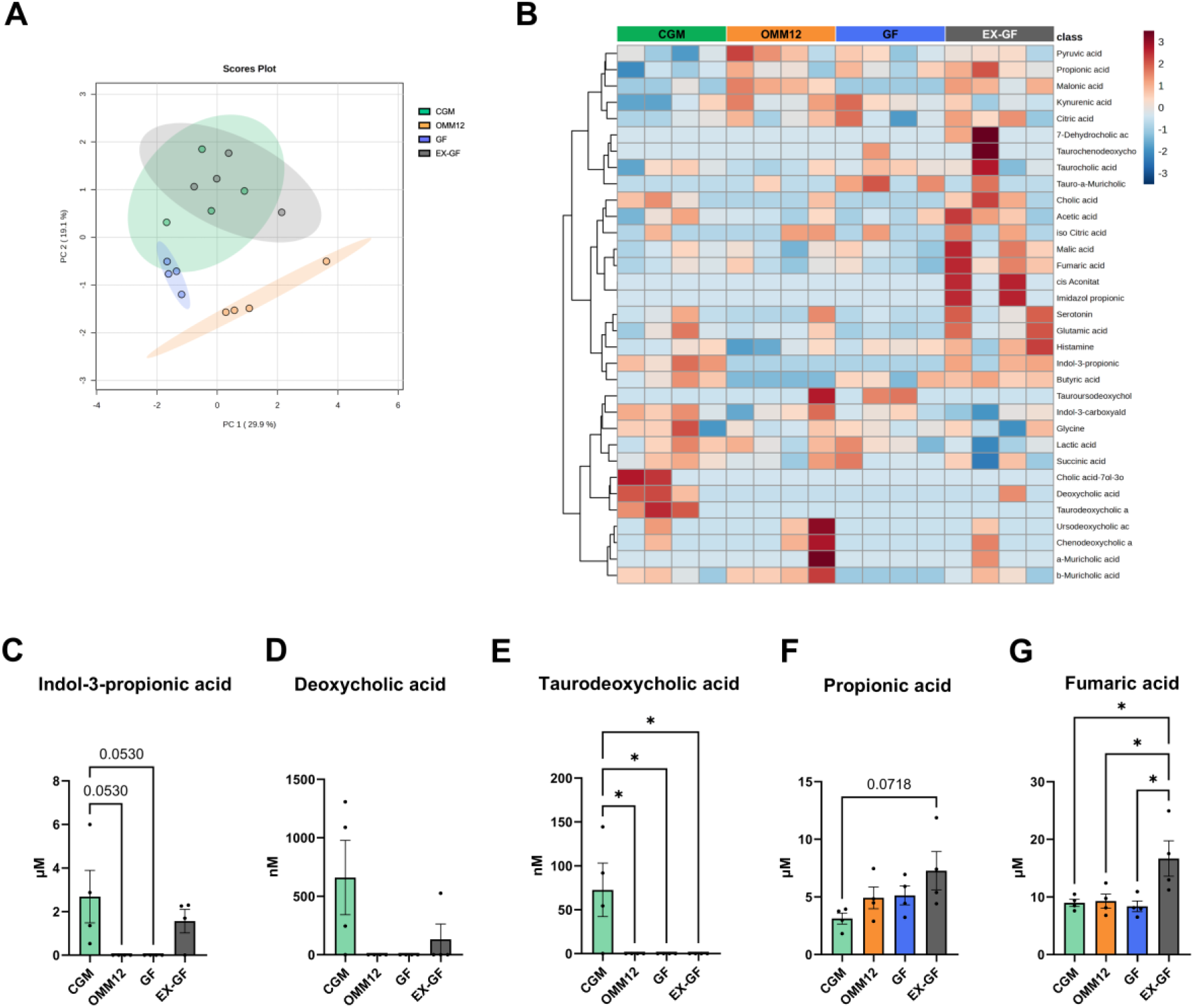
Analysis of gut microbiota-associated serum metabolites. (A) Principal component analysis (PCA) scores plot showing the distribution of individual samples based on metabolite profiles across experimental groups. (B) Heatmap displaying the relative abundance of targeted metabolites, including short-chain fatty acids (SCFAs), bile acids, and neurotransmitter-related compounds. Each row represents a specific metabolite, and each column corresponds to an individual sample. Color intensity reflects relative metabolite levels, with higher concentrations shown in orange to red and lower concentrations in light to dark blue. (C–G) Serum concentrations of selected gut-derived metabolites. One-way ANOVA for parametric data, or Kruskal-Wallis for non-parametric data; *p ≤ 0.05. n=4 mice, each group. Data are presented as mean ± SEM.

Among all detected metabolites, including short-chain fatty acids (SCFAs), bile acids, and neurotransmitters (Fig. 10B), five showed differential abundance among groups with different recovery patterns in EX-GF samples (Fig. 10 C-G). In particular, indole-3- propionic acid (IPA), undetectable in GF and OMM12 mice, was restored following post-weaning microbiota colonization (Fig. 10C). In contrast, the secondary bile acids deoxycholic acid (DCA) and taurodeoxycholic acid (TDCA) remained largely absent in EX-GF mice (Fig. 10D, E), indicating incomplete restoration of specific microbiota-dependent metabolic pathways. Serum levels of propionic and fumaric acid, unchanged among all other groups, were increased following post-weaning microbiota colonization (Fig. 10F, G).

Together, these findings highlight distinct microbial and metabolic signatures associated with differential neuromuscular recovery following post-weaning microbiota colonization.

## Discussion

The gut microbiota has emerged as an important regulator of neuromuscular development (7, 8) In our previous work, we demonstrated that the absence of a microbiota impairs the normal development of peripheral nerves, skeletal muscles and neuromuscular junctions. Furthermore, we showed that these alterations were not completely reversed in gnotobiotic animals colonized with the simplified OMM12 microbial consortium (7, 8). This prompted us to investigate whether these developmental alterations could be reversed by restoration of a more complex microbiota. To address this issue, we colonized germ-free mice by co-housing them with conventionally raised animals immediately after weaning and examined peripheral nerves, skeletal muscles and neuromuscular junctions at the adult stage. Here, we show that, although this approach successfully transferred a complex GM in EX-GF mice, its effects were clearly different across tissues, with skeletal muscle showing extensive structural and molecular recovery, opposite to peripheral nerves, showing persisting abnormalities. Together, these findings indicate that microbiota absence-dependent developmental alterations are not equally reversible and that the outcome of post-weaning microbiota restoration differs substantially between skeletal muscle and peripheral nerves.

Co-housing efficiently introduced a complex gut microbiota to EX-GF mice, likely through environmental microbial exposure and coprophagy (13). As a result, EX-GF mice developed a microbial community broadly resembling that of conventionally raised animals. However, differences in the relative abundance of several bacterial taxa indicated that the constituted microbiota did not fully replicate the donor community. These compositional differences were accompanied by incomplete recovery of selected circulating microbial metabolites. Although a complex microbiota was successfully established, the persistent differences in microbial composition and metabolic profile may have contributed to the distinct recovery patterns observed across the neuromuscular system.

The persistence of peripheral nerve abnormalities following post-weaning GM colonization was unexpected. Despite successful microbial transfer, nerve structural alterations remained evident and were accompanied by further transcriptional changes. This indicates that delayed restoration of the microbiota is insufficient to fully normalize peripheral nerve development. Indeed, in rodents, myelination progresses rapidly during the first three postnatal weeks (14–16), but notably throughout this period both GF and EX-GF mice developed in the absence of microbiota. We believe that microbiota-derived signals that might be required and act during this key developmental period were missing, and GM colonization after weaning failed in recovering myelination, because pivotal developmental programs had already taken place. One explanation for these findings may lie in the biology of Schwann cells. During development, Schwann cells progressively establish stable myelin sheaths around axons, a process that is largely completed during the early postnatal period (14–16). In the absence of nerve injury, mature Schwann cells do not normally disengage from intact axons to rebuild existing myelin. If microbiota-derived signals contribute to this developmental program, their absence during early postnatal life may have lasting consequences that cannot be fully corrected by delayed colonization. A comparable timing-dependent phenomenon has been reported in the central nervous system, where GM depletion leads to irreversible deficits (17–19). Although our data are consistent with this view, further studies are required to enable us to distinguish whether the limited recovery reflects the timing of microbial exposure, the absence of specific microbiota-derived signals during this developmental window, or a combination of both.

The molecular changes observed in EX-GF nerves are consistent with persistent alterations in axon-Schwann cell interactions. Ultrastructural analysis revealed that hypermyelination was caused by excessive Schwann cell wrapping, as evidenced by the increased number of myelin lamellae. This phenotype was accompanied by elevated expression of axonal type III Neuregulin-1 (*Nrg1*), a key axonal regulator of Schwann cell myelination and myelin thickness (11, 20). Interestingly, hypermyelination occurred without the upregulation of core myelin genes, but was accompanied by increased expression of peripheral myelin protein 2 (*Pmp2*), a lipid-binding protein found in compact myelin. Previous studies have shown that overexpression of NRG1 type III promotes hypermyelination and induces *Pmp2* expression in experimental models of congenital hypomyelination and Charcot-Marie-Tooth1B (20, 21). PMP2 is a downstream target of NRG1 signaling and facilitates fatty acid uptake by myelinated nerve fibers (22), suggesting that activation of the NRG1-PMP2 axis may support both excessive myelin wrapping and the associated metabolic demands. It is unclear whether activation of this pathway contributes directly to the observed phenotype, represents a compensatory response to abnormal myelin development or reflects another consequence of altered microbiota-dependent development. Similarly, the elongation of nodes of Ranvier, together with the dysregulation of node-associated genes, further supports the persistence of altered axon-glia interactions following post-weaning microbiota colonization.

The most striking finding of this study is that skeletal muscle responded very differently to delayed microbiota restoration. Unlike peripheral nerves, both the structural and molecular features of skeletal muscle were largely normalized following post-weaning colonization. This is consistent with previous studies showing that muscle loss caused by the absence or depletion of the gut microbiota - whether in germ-free animals or following antibiotic treatment - is reversible upon microbiota (re)colonization (8, 23, 24). Here, we extend these observations by showing that recovery is not limited to muscle mass, but also involves normalization of protein homeostasis, broad proteomic remodeling, metabolic pathways and sarcomere organization. In addition, comparison of two anatomically and functionally distinct hindlimb muscles revealed that the effects of microbiota depletion are not uniform. While both TA and GA muscles recovered following post-weaning colonization, the simplified OMM12 consortium resembled the TA phenotype but only partially resembled the GA phenotype of CGM mice. This observation is consistent with previous studies showing that individual hindlimb muscles differ in their susceptibility to disease and ageing (25–27), suggesting that the recovery of individual skeletal muscles may depend on the composition of the newly established microbial community and the repertoire of metabolites it produces. Together, these findings indicate that the effects of microbiota absence on skeletal muscle are largely reversible under the experimental EX-GF conditions examined here.

The proteomic analysis provides a broader perspective on the recovery of skeletal muscle following post-weaning GM colonization and, to our knowledge, represents the first comprehensive characterization of skeletal muscle proteomes across different microbiota conditions. Rather than affecting a limited number of pathways, the absence of a complex microbiota reshaped multiple aspects of muscle biology, including energy metabolism, protein turnover and cytoskeletal organization. Most of these molecular changes were no longer evident in EX-GF muscles, indicating that post-weaning GM colonization is sufficient to restore a proteomic profile closely resembling that of conventionally raised animals. These findings extend previous observations linking the gut microbiota to muscle metabolism and fiber composition (8, 9, 28), while revealing sarcomere-associated proteins as an additional component of microbiota-dependent muscle remodeling. This molecular recovery was mirrored at the ultrastructural level. Although overall sarcomere architecture remained preserved, GF muscles exhibited increased Z-line misalignment and a higher frequency of structural abnormalities, both of which were largely corrected following post-weaning colonization. Since disruption of Z-line organization is a hallmark of several myopathies and is associated with impaired muscle integrity (29–31), these observations suggest that the impact of the gut microbiota extends beyond metabolic regulation to the structural organization of the contractile apparatus. To our knowledge, this is the first study to identify structural modeling of the sarcomere as a feature of microbiota-dependent skeletal muscle development. Although the influence of the gut microbiota on muscle metabolism and fiber composition is now well established, differences in experimental models, the muscles analyzed, and the strategies used to manipulate and restore the microbiota have contributed to variable findings across studies [8,9,55]. Our data suggest that at least part of this variability may reflect differences in the repertoire of microbiota-derived metabolites available to skeletal muscle [8,9].

The neuromuscular junction further illustrates the differential reversibility observed across the neuromuscular system. While postsynaptic fragmentation was largely restored following post-weaning colonization, several presynaptic abnormalities persisted, mirroring the incomplete rescue observed in peripheral nerves. This dissociation between pre- and postsynaptic compartments is consistent with the broader divergence between skeletal muscle and peripheral nerve phenotypes. Despite the persistence of structural and molecular abnormalities in peripheral nerves in EX-GF mice, skeletal muscle showed extensive rescue following post-weaning microbiota constitution. Similar dissociation between pre- and postsynaptic alterations has also been described in other neuromuscular disorders (32). Although these findings suggest that physiologic skeletal muscle phenotypes can develop despite persistent nerve abnormalities, neuromuscular performance and muscle force were not assessed in the present study. Therefore, whether the residual presynaptic alterations have functional consequences for neuromuscular transmission or muscle performance remains to be determined.

The distinct patterns of rescue observed across the neuromuscular system were accompanied by selective resembling of circulating microbiota-derived metabolites. Among these, IPA appeared after post-weaning colonization, whereas secondary bile acids such as DCA and TDCA remained largely absent. These findings indicate that successful microbial colonization does not necessarily result in complete constitution of microbiota-dependent metabolic functions, reinforcing the concept that microbial composition and metabolic output establish at different rates.

IPA has attracted considerable interest because of its reported effects on skeletal muscle and peripheral nerves (33–35). In our study, rescue of circulating IPA levels coincided with the rescue of skeletal muscle but not of peripheral nerve abnormalities. These observations do not support a direct relationship between circulating IPA and the rescue of the nerve phenotype. At the same time, they do not exclude the possibility that IPA, or other microbiota-derived metabolites, might contribute to peripheral nerve development if present during an earlier stage of postnatal maturation. Addressing this question will require experiments specifically designed to manipulate metabolite availability during defined developmental windows.

Among the bacterial taxa enriched following colonization, *Duncaniella muris* strain B8 deserves particular attention. Recent work identified this species as a contributor to the restoration of circulating IPA levels after stroke (36), making it an interesting candidate for future investigations able to better define a causal relationship between the expansion of *Duncaniella*, IPA availability and the phenotypic recovery observed in EX-GF mice.

## Conclusion

Our findings show that microbiota-dependent developmental abnormalities differ in their reversibility after post-weaning colonization. Establishing a complex gut microbiota after weaning broadly normalized skeletal muscle structure and molecular programs but did not reverse peripheral nerve abnormalities. Whether this difference reflects the timing of microbial exposure, incomplete restoration of specific microbiota-derived signals, or both remains unresolved. Testing earlier (or later) colonization and temporally controlled metabolite supplementation will be important for defining the developmental requirements of the gut-peripheral nerve axis.

## Materials and Methods

### Mice

A total of 41 63-76-day-old adult mice with different hygiene status, grown in the Central Animal Facility (Hannover Medical School, Hannover, Germany) were used for this study. As previously described (7), the hygiene status was: specific pathogen-free C57BL6/JZtm (harboring complex gut microbiota/CGM), gnotobiotic (colonized with Oligo-Mouse-Microbiota 12, OMM12), germ-free (GF), and ex-germ-free (EX-GF). To generate CGM and OMM12 mice, GF C57BL6/JZtm mice were colonized with their respective microbiota. The formation of substrains was avoided by repeating this procedure every 10 generations. GF and gnotobiotic OMM12 mice were bred and maintained in isolators (Metall+Plastik GmbH, Radolfzell-Stahringen, Germany) placed in a controlled environment (50–55% humidity, 20–22°C,14:10-hour light/dark cycles). EX-GF mice were generated by co-housing GF off-spring together with CGM off-spring from weaning onward. GF and gnotobiotic OMM12 mice were fed as previously described in detail (7). Specific pathogen-free mice and co-housed CGM and EX-GF mice were housed in individually ventilated cages (XJ Edge, Allentown) in the same controlled environmental conditions for humidity, temperature and light/dark cycle. Specific pathogen-free and EX-GF mice were fed with 50 kGy gamma-irradiated diet (#1314, Altromin) and water at libitum, as previously described (7). The germ-free (GF) and gnotobiotic mouse colonies are subjected to extensive and rigorous testing to verify their microbiological status. The Central Animal Facility at the Medical School Hannover (MHH, Hannover, Germany) follows a comprehensive set of screening procedures that combine classical microbiology with molecular-biological techniques. Mold traps are kept in the isolators at all times to detect any incursion of molds or yeasts. On a monthly basis, fecal pellets, environmental samples (e.g., feed, bedding) and swabs are taken from the isolators and examined for bacterial and fungal contamination. Samples are inoculated directly into thioglycolate broth (for detection of aerobes, anaerobes, microaerophiles and fastidious organisms) and Sabouraud dextrose broth (for detection of yeasts, molds and dermatophytes) and incubated under aerobic conditions at 37 °C, 29 °C and ambient temperature for at least 10 days. Every six months, retired breeder mice are euthanized for a detailed assessment of specific mouse pathogens in accordance with the FELASA recommendation list (37) and to confirm the gnotobiotic status of the colonies. For this, cultures of cecal and colonic contents, direct phase-contrast microscopy of native cecal smears, and evaluation of cecum size (shrunk cecum in GF animals could indicate microbial contamination). OMM12 breeding colonies are additionally screened by 16S-rRNA-based, microbe-specific qPCR and by next-generation sequencing. These methods confirm the presence of the intentionally introduced microorganisms and exclude unknown or unwanted contaminants (38). With this we ensure that mice used in our study have not revealed any infections with common murine pathogens in CGM animals, nor any contaminations in the GF or OMM12 mice, so that the latter two remained free of undesired microorganisms.

Animals were euthanized by carbon dioxide (CO₂) inhalation. CO₂ exposure was used to induce unconsciousness, followed by complete respiratory arrest. Death was confirmed prior to any further procedures. The study complied with the German Animal Welfare Legislation and the principles of the Basel Declaration and recommendations of Directive 2010/63/EU. All procedures received the approval by the local Institutional Animal Care and Research Advisory Committee and were registered with the Animal Care Committee of Lower-Saxony, Germany. The registration number for breeding and mouse lines is 42500/1H according to §11 of the German protection of animal act (TierSchG). Announcement of animal sacrifice for scientific purposes was given to the authorities under the numbers §4 2017-171, §4 2022-308.

### DNA isolation, shallow shotgun metagenomics and sequencing bioinformatic analysis

Fecal pellets from n=4 CGM and n=5 EX-GF mice were collected, stored at -80°C and sent to “Novogene” for DNA isolation and shotgun metagenomics for taxonomical profiling of the gut microbiota.

### BCL to FASTQ conversion and quality control

BCL files were converted to FASTQ files using bcl2fastq Conversion Software version v2.20.0.422 (Illumina). Sequencing read quality control was performed with FastQC (Andrews, S. (2010). FastQC: A Quality Control Tool for High Throughput Sequence Data [Online]. Available online at: https://www.bioinformatics.babraham.ac.uk/projects/fastqc/). The data integration and summarizing visualization was created with MultiQC software (39).

### Taxonomic classification of sequencing reads

KrakenUniq metagenome analysis pipelines with default settings were applied to determine hypotheses regarding the origin of the sequencing reads of the metagenomic samples. KrakenUniq (REF Salzberg, Version 1.0.4), has been well described already elsewhere. The “MicrobialDB” reference database, available from https://benlangmead.github.io/aws-indexes/k2 -> MicrobialDB (2023-08-08; https://genome-idx.s3.amazonaws.com/kraken/uniq/krakendb-2023-08-08-MICROBIAL/kuniq_microbialdb_minus_kdb.20230808.tgz) was used for the analysis.

The Bracken framework was used to calculate the frequency of species in DNA sequences from the metagenomics samples obtained (https://github.com/jenniferlu717/Bracken, Bracken v2.9; Bracken’s peer-reviewed paper (published Jan 2, 2017): “Bracken: estimating species abundance in metagenomics data” https://peerj.com/articles/cs-104/). Alpha and beta diversity were calculated using the implementation of the KrakenTools suite (40).

The pcoa implementation of the ’ape’ R package (ape version 5.8 (41)) was used to visualize beta diversity using Principal Coordinate Analysis (PCoA).

### RNA extraction and RNA-seq library preparation and analysis

For RNA-seq, sciatic nerves from each animal were pooled. For dorsal root ganglia (DRG), ∼8–10 randomly collected DRG along the spinal column were pooled. RNA-seq was performed on samples from three animals per experimental group, providing three independent biological replicates per group. Total RNA was isolated using TRIzol reagent (Life Technologies) and processed following the manufacturer’s instructions. RNAseq data from CGM, OMM12 and GF groups were obtained from previous work (7). RNA-seq data from DRG and sciatic nerves EX-GF samples were processed consistently with the previous work: good-quality reads were aligned to the mouse reference genome (UCSC mm10) using STAR 2.7.1a (42) (-- outFilterMismatchNmax 999 -- outFilterMismatchNoverLmax 0.04) and raw counts were quantified with featureCounts v1.6.3 (43) (-t exon -g gene_name) using the GENCODE M23 annotation. Raw counts from CGM, OMM12, and GF groups were obtained from previous work and integrated. To assess potential batch effects, PCA was performed on normalized expression values across all samples. Raw counts were then separated by tissue of origin, and genes with low expression across samples were filtered prior to normalization (1 CPM in fewer than 3 samples). Library size differences were corrected using TMM normalization, and expression values were log2-transformed CPMs with the EdgeR package (44). Differential expression analysis was performed by fitting a GLM to all groups and performing the LF test for pairwise comparisons with the formula “∼0+condition”. Genes were considered DE according to precedent thresholds (abs. logFC > 1 and p-value < 0.01). Gene set enrichment analysis was conducted using GSEA software (45) on the log2FC*(-logPval) ranked genes. Gene expression heatmaps were generated using the ComplexHeatmap R package, with logCPM values scaled to Z-scores across samples (46).

### Analysis of histological preparations and electron microscopy images of nerves

As in our previous study (7), we analyzed median nerve histomorphometry to assess myelinated fiber changes. Compared to our earlier data (N=7 per group), we increased sample sizes to 9–11 per group for CGM, OMM12, GF and included the EX-GF group.

Median nerves were collected and analyzed according to an established protocol [20, 21]. In brief, Karnovsky solution (2% paraformaldehyde, 2.5% glutaraldehyde in 0.2 M sodium cacodylate buffer, pH 7.3) for 24 h was applied to fix 5 mm long nerve samples. Samples were then washed in 0.1 M sodium cacodylate, 7.5% sucrose and further fixed (1% osmium tetroxide for 1.5 h).

Stereological analysis was performed as previously described (47, 48) using light microscopy (Olympus Europa SE & Co. KG, Hamburg, Germany) equipped with a prior controller and Stereo Investigator software, version 2021.1.1 (MBF Bioscience, Williston, VT, USA). From each sample, in two randomly selected sections optical fractionator was used to determine cross-sectional area (20x magnification), number of myelinated fibres, axon diameter, fiber diameter, myelin thickness (100x magnification), and to calculate the nerve fiber density and g-ratio (version 1.48; National Institutes of Health, Bethesda, USA). For ultrastructural analysis, resin-embedded samples were sectioned into ultrathin 70 nm thick slices and imaged using a JEM-1010 transmission electron microscope (JEOL, Tokyo, Japan). Specifically, the length of the nodes of Ranvier and myelin lamellae were analyzed using ImageJ software.

### Quantitative Real Time PCR (qRT-PCR)

RNA extraction, reverse transcription and qRT-PCR analysis of gastrocnemius (GA) muscle samples were performed as previously described (7). Data were normalized to Ribosomal Protein L7 (*Rpl7l1*) expression levels and to the mean value of the control group. The following primer sequences were used: *Fbxo32*: forward: 5’– TCGACTGCCATCCTGGATTC –3’, reverse: 5’– TTCTTTTGGGCGATGCCACT –3’; *Trim63*: forward: 5’– ACCTGCTGGTGGAAAACATC –3’, reverse: 5’– CTTCGTGTTCCTTGCACATC –3’; *Myh1*: forward: 5’– TTCATTAGTTTCCCAGCTCTCC –3’, reverse: 5’– AGGCACTCTTGGCCTTTATC –3’; *Myh2*: forward: 5’– GGCTTCAGGATTTGGTGGATAA –3’, reverse: 5’– GGATCTTGCGGAACTTGGATAG – 3’; *Myh4*: forward: 5’– ATTGACGTGGAGAGGTCTAAC –3’, reverse: 5’– CCTGAGTTTCCTCGTACTTCTG –3’; *Myh7*: forward: 5’–, CCATCTCTGACAACGCCTATC –3’, reverse: 5’– GGATGACCCTCTTAGTGTTGAC –3’; *Rpl7l1*: forward: 5’– AGAGCAGGAGCAGGTTTTCC –3’, reverse: 5’– CAGCCAATGAGGGAACTCGT –3’.

DRG samples were processed following a protocol previously described (7). Data were normalized to Tbp expression and to the mean value of the control group. The following primer sequences were used: *Nrg1-Type III*: forward: 5’– CCCTGAGGTGAGAACACCCAAGTC –3’, reverse: 5’– TGGTCCCAGTCGTGGATGTAGATG –3’; *Tbp*: forward: 5’– GATCAAACCCAGAATTGTTCTCC –3’, reverse: 5’– GGGGTAGATGTTTTCAAATGCTTC –3’.

### Whole-mount staining and NMJ morphometric analysis

Hemidiaphragms were fixed with 4% PFA in PBS for 1 h at room temperature. After removing the surrounding connective tissue, muscles were washed in PBS supplemented with 0.2% Triton-X-100 detergent, saturated with a blocking solution [10 % goat serum (GS), 2% Triton-X-100 in TBS] overnight at 4°C, and then incubated for 48 h at 4°C with the following primary antibodies diluted in a solution of TBS containing 1% GS and 2% Triton-X-100: rabbit anti-peripherin (1:200, Novus Biologicals, NB300-137) and rabbit anti-synaptophysin (1:50, Santa Cruz Biotechnology, sc-9116). Diaphragms were extensively washed in TBS and stained overnight at 4°C with Alexa Fluor™ 555 conjugated α- bungarotoxin (αBT) (1:1000, Invitrogen) and anti-rabbit Cy2 (1:200, Jackson Immunoresearch), diluted in TBS supplemented with 2% GS, 0.02% Triton X-100. After washing, samples were mounted in 80% glycerol-PBS and imaged using a Leica Stellaris SP8 confocal microscope. Z-stack images of at least 40 en face NMJs per sample were acquired with the following settings: 63x objective, 512 x 512 frame size, 400 speed, 2.5x zoom, 2 line average and 1 μm z-stack interval. Maximum intensity Z-stack projections were analysed with the aNMJ-morph macro of the Fiji software to derive morphometric features of NMJs, including the following pre- and postsynaptic parameters: AChR area, AChR perimeter, endplate area, endplate perimeter, compactness [(AChRarea/endplate area) x100], axon diameter, nerve terminal perimeter, nerve terminal area, and complexity [log10(n. terminal branches x n. branch points x total length of branches)] (12). Finally, the degree of overlap between pre- and postsynaptic elements was measured [(total area AChRs - unoccupied area AChRs)/total area AChRs x 100]. The number of AChRs clusters per NMJ was determined by visual inspection of Z-stack images using the 3D view in the AIVIA AI image analysis software and it was used to derive the fragmentation index [1-(1/number of AChR clusters)], where a value of zero reflects intact, plaque-like NMJs as opposed to a value tending to one, indicative of highly fragmented endplates.

### Transmission electron microscopy of muscle samples

Fresh tissue samples of diaphragm muscles were fixed in Karnovsky solution (2% paraformaldehyde, 2.5% glutaraldehyde in 0.2 M sodium cacodylate buffer, pH 7.3) for 24 h, washed in 0.1 M sodium cacodylate buffer, post–fixed for 1.5 h in 1% osmium tetroxide and embedded in epoxide resin. Ultrathin sections were cut to visualize the longitudinal axis of the myofibrils and stained with uranyl acetate and lead citrate. Images were acquired using a FEI Tecnai G2 transmission electron microscope (Electron Microscopy Service, Biology Department, University of Padova) equipped with an Olympus Veleta CCD digital camera (Olympus Soft Imaging System). Sarcomeric morphological parameters, including sarcomere length and height, A-band and I-band length, Z-line length and width, were measured manually using Fiji software in at least 80 sarcomeres per muscle, sampled from four to nine different images. To quantify Z-line misalignment, the displacement of Z-lines in adjacent myofibrils was measured using the Fiji software and the multi-point tool as the lateral distance between the midpoint of two consecutive Z-lines. Displacements were measured along continuous rows of Z-lines (Z-staircases) and averaged across multiple Z-staircases. Phenotypic appearance of Z-lines was assessed by visual inspection. At least 12 Z-staircases and 150 Z-lines were analysed for each muscle, sampled from four to nine separate locations. All analyses were conducted in blind.

### Protein extraction and western blot analysis of muscle samples

Total protein lysates were extracted from mechanically pulverized frozen Tibialis anterior (TA) and GA muscles and analysed through Western blot analysis as previously described [7]. The following primary antibodies were used: anti-LC3B (raised in rabbit, 1:1000, PA1- 16930, Thermo Fisher Scientific,); anti-phospho-S6 Ribosomal Protein (Ser240/244) (raised in rabbit, 1:1000, Cell Signaling Technology, #5364); anti-S6 Ribosomal Protein (raised in rabbit, 1:1000, Cell Signaling Technology, #2217), anti-vinculin (raised in mouse, 1:1000, Sigma-Aldrich, V4505). Fiji software was used for densitometric quantification. Phosphorylated forms were normalized to the corresponding total protein pair loaded on the same membrane.

### Protein Extraction and Label-Free Liquid Chromatography with Tandem Mass Spectrometry

GA muscle biopsies were pulverized, suspended in lysis buffer and processed to quantification as previously described (49). 100 µg per sample of protein extracts were processed after the FASP protocol as described (50). Each sample was deposited in a Microcon-30 kDa centrifugal filter unit (Merck Millipore, Burlington, MA, USA) and washed by centrifugation at 14,000 g for 15 min with 200 µL of UA buffer (8 M urea, 0.1 M Tris/HCl, pH 8.5). Samples were carbamidomethylated in 100 µL of 50 mM iodoacetamide in UA buffer for 20 min, then washed three times in 100 µL UA buffer followed by three washes in 100 µL of 50 mM ammonium bicarbonate in water. Filters were incubated with sequence grade trypsin (Promega, Madison, WI, USA) for 16 h at 37 °C using a protein:trypsin ratio of 50:1. After acidification with trifluoracetic acid and desalting on C18 tips (Zip-Tip C18 micro, Merck Millipore, Burlington, MA, USA), peptide samples were vacuum concentrated, reconstituted in HPLC buffer A (0.1% formic acid) and separated with an Easy Spray PepMap RSLC C18 column (25 cm, int. diameter of 75 µm,Thermo Fisher Scientific), on a Dionex UltiMate 3000 HPLC System, connected to an Orbitrap Fusion Tribrid mass spectrometer (Thermo Fisher Scientific). For each sample, three technical replicates were analyzed. Mass spectra were processed using MaxQuant software (Max Planck Institute of Biochemistry, Munich, Germany, version 1.6.17.0) (51), allowing a maximal mass deviation of 6 ppm for monoisotopic precursor ions and of 0.5 Da for MS/MS peaks. Further criteria included enzyme specificity, set to trypsin/P, with two missed cleavages allowed as a maximum, fixed modification, set as carbamidomethylation and variable modifications as N-terminal acetylation and methionine oxidation. Andromeda search engine was applied against the Mus musculus Uniprot UP000000589 sequence database (54707 proteins, March 2024). At least one unique or razor peptide per protein group was required for protein identification. MaxQuant quantification exploited the built-in XIC-based label-free quantification algorithm using fast quantification (52). FDR was set to 1% at the peptide, at the protein, and 1at the site-modification level, and 7 amino acids was set as the minimum required peptide length. Perseus software was used to assess statistical analyses (v.1.6.15.0, Max Planck Institute of Biochemistry, Martinsried, Germany) (53). Only proteins identified in at least 80% of samples were considered, for each experimental group. The Benjamini–Hochberg false discovery rate test was applied to reduce false positives.

### Ingenuity Pathway Analysis and annotation

Functional annotation of differentially regulated proteins was curated manually combining information from KEGG, GO, and Uniprot databases as well as literature reports. The software Ingenuity Pathway Analysis (IPA, winter release 2023; Qiagen, Hilden, Germany) was used for functional and network analyses of protein expression changes as previously described (49). Biological pathways enriched with differentially regulated proteins were assessed, by applying the “core analysis” function. Significant proteins or regulators across experimental conditions were then visualized and identified by applying the “comparison analysis” function. p-values were assessed by a right-tailed Fisher’s exact test. Activation/inhibition of a pathway/regulator/disease and biofunction were predicted using the activation z-score (54). Ap-value < 0.05 and z-scores ≤ −2 and ≥2, which considers the directionality of the observed effects, were considered statistically significant by applying Fisher’s exact test.

### Teased Nerve Fiber Preparation and Internodal Length Measurement

Three ulnar nerves from each experimental group were fixed in 4% paraformaldehyde for 24 hours, followed by impregnation with 1% osmium tetroxide in 0.1 M Sorensen buffer. Afterward, the samples were washed twice in 70% ethanol to remove excess osmium and then transferred to glycerin. Individual nerve fibers were carefully separated using fine-tipped tweezers under a stereomicroscope and mounted on glass slides in glycerin to prevent dehydration. Nodes of Ranvier were identified under a light microscope, and images of intact internodal segments (between two consecutive nodes) were captured using a calibrated digital camera. Internodal lengths were measured with the image analysis software “IM50”.

### Histological staining and morphometric analysis on tibialis anterior (TA) muscles

4% paraformaldehyde (PFA)-fixed TA muscles were dehydrated in ethanol, cleared with toluene and paraffin embedded as previously described (7). 7-μm-thick transversal sections were dewaxed with xylene, and rehydrated in ethanol before labeling with Alexa Fluor™ 488 conjugated Wheat Germ Agglutinin (WGA, 2 μg/mL, Invitrogen) and Hoechst 33342 (1 μg/mL, Invitrogen) in PBS at room temperature for 20 min. After three washes in PBS, slides were mounted in 80% glycerol in PBS. For each sample, images of entire muscle sections were acquired at a 10x magnification using a Leica Stellaris SP8 confocal microscope with the Leica Application Suite X (LASX) software in the Navigator acquisition mode. Images were segmented using the Cellpose object detection recipe in the AIVIA AI image analysis software. Wrong selections were manually corrected. Cross-sectional area (CSA) and minimum Feret’s diameter (MFD) measurements were obtained from AIVIA- generated masks using the LabelsToRois plugin of Fiji (55).

### Immunofluorescence staining and morphometric analysis on gastrocnemius muscles (GA)

10-μm-thick GA cryosections were blocked for 1 h with 4% IgG-free Bovine Serum Albumin (BSA, Sigma) in PBS, followed by 1h incubation with anti-mouse IgG Fab fragment (1:25, Jackson Immunoresearch) in 4% IgG-free BSA in PBS, and then immunostained with the following primary antibodies, diluted in 0.5% IgG-free BSA in PBS: mouse anti-myosin heavy chain (MHC)-I (1:25, DSHB BA-D5), mouse anti-MHC-IIA (1:50, DSHB SC-71), mouse anti-MHC-IIB (1:50, DSHB BF-F3), and rabbit anti-laminin (1:100, kindly supplied by G. M. Bressan, Padova). After washing in PBS, sections were stained with proper secondary antibodies: anti-mouse Cy3 (1:250, Jackson Immunoresearch), anti-mouse IgG1 Alexa Fluor™ 568 (1:250, Invitrogen), anti-mouse IgM Alexa Fluor™ 488 (1:250, Invitrogen), anti-rabbit Cy2 (1:250, Jackson Immunoresearch) and anti-rabbit Alexa Fluor™ 647 (1:100, Invitrogen). Slides were mounted with the Fluoroshield histology mounting medium (Sigma-Aldrich, F6182). Entire muscle sections were imaged at a 10x magnification using a Leica Stellaris SP8 confocal microscope coupled with the Navigator acquisition mode of the LASX software. Morphometric and fiber type analysis was performed using the AIVIA AI imaging analysis software. Images were first segmented using the Cellpose object detection recipe and then fibers belonging to each fiber subtype were identified using the Object Classifier deep learning tool. The LabelsToRois plugin of ImageJ was used to derive CSA and MFD measurements related to all fibers and to each fiber type subgroup (55).

### Metabolomics

#### Sample preparation of mouse serum

50 µL of Serum was extracted with 250 µL of methanol-based dehydrocholic acid extraction solvent (c=1.3 µmol/L). The samples were extracted for 15 min at 10°C at 1000 rpm using an Eppendorf Thermomix (Eppendorf, Hamburg, Germany). Afterwards the samples were centrifuged for 10 min at 10°C at 15.000 rpm (Eppendorf Centrifuge 5424R, Hamburg, Germany).

#### Targeted bile acid measurement

A volume of 20 µL of isotopically labeled bile acids (ca. 7 µM each) was added to 100 µL of sample extract. QTRAP 5500 triple quadrupole mass spectrometer (Sciex, Darmstadt, Germany) integrated with an ExionLC AD (Sciex, Darmstadt, Germany) ultrahigh performance liquid chromatography system was used to make a quantification of the targeted bile acids. Detection and quantification were performed using multiple reaction monitoring (MRM) as described by Reiter et. al. (56, 57). Data collection and instrumental control were performed with Analyst 1.7 software (Sciex, Darmstadt, Germany).

#### Targeted neurotransmitter measurement

The same sample extract was used for the neurotransmitter measurement. The MRM-transitions were optimized for adrenaline, noradrenaline, acetylcholine, gamma-amino butyric acid, serotonin, histamine, dopamine, glycine, glutamic acid, cortisol, kynurenic acid, 4-hydroxy-3-methoxymandelic acid, homovanillic acid. The same LC-MS-set up as mentioned above was used. A Premier BEH Amide 2.1 x 100 mm, 1.7 µm column (Waters, Eschborn, Germany) was used for chromatographic separation. Solvent A was 30 mM ammonium formate in water/acetonitrile (95/5, v/v) at pH 3 and solvent B was 30 mM ammonium formate in water/acetonitrile (15/85, v/v) at pH 3. The gradient started with 100% B for 2 min. At 3.5 min the gradient was decreased to 35% B which was held until 6.5 min. Afterward the gradient was increased again to 100% B again which was held for 3 min. The column oven was set to 30°C, the flow rate was 400 µl/min and the injection volume was 5 µL. The electrospray voltage was set to 5500 V, curtain gas to 40 psi, ion source gas 1 to 55 psi and ion source gas 2 to 65 psi. Data acquisition and instrumental control were performed with Analyst 1.7 software (Sciex, Darmstadt, Germany).

#### Targeted short chain fatty acid (SCFA), organic acid and tryptophane metabolite measurement

The 3-NPH method was used for the quantification of SCFAs (58, 59). Briefly, 80 µL of the serum extract and 30 µL of isotopically labeled standards (ca 50 µM) were mixed with 20 µL 120 mM EDC HCl-6% pyridine-solution and 20 µL of 200 mM 3-NPH HCL solution. After 30 min at 40°C and shaking at 1000 rpm using an Eppendorf Thermomix (Eppendorf, Hamburg, Germany), 850 µL acetonitrile/water (50/50, v/v) was added. After centrifugation at 13000 U/min for 2 min the clear supernatant was used for analysis. The same system as described above was used. A multiple reaction monitoring (MRM) method was used for the detection and quantification of SCFAs, organic acids and tryptophane metabolites according to Hemmati et al. (60).

#### Statistical Analysis

GraphPad Prism Version 10.2.3 or 10.6.1 (GraphPad Software, San Diego, CA, USA) was used for statistical analyses. Data are presented as mean ± SEM. Outliers were detected using Grubb’s test and excluded from subsequent analysis. Normal distribution was assessed by applying the Shapiro-Wilk test. One-way ANOVA followed by Tukey’s multiple comparisons post-hoc test, unpaired two-tailed Student’s t test or two-way ANOVA test followed by uncorrected Fisher’s LSD test were applied to evaluate statistical significance for parametric data, as indicated. Non-parametric data were analyzed through the Kruskal-Wallis test followed by Dunn’s multiple comparisons post-hoc test. Statistical significance was set at a p-value ≤ 0.05. N corresponds to the number of mice investigated per group, except where otherwise stated.

## Acknowledgments

We thank Jennifer Metzen, Silke Fischer, Natascha Heidrich, and Melanie Weiß (Institute of Neuroanatomy and Cell Biology, Hannover Medical School, Germany), Anna Smoczek, Tim Scheele, Jasmin Goralski and animal caretakers in the Gnotobiotic unit (Institute for Laboratory Animals Science) for their excellent technical assistance.

## Data and materials availability

All data required to support the conclusions of this study are included in the article and its Supplementary Materials. All datasets generated during this study have been deposited in appropriate repositories.

RNA-seq datasets for CGM, GF, and OMM12 samples were obtained from our previously published study (7) and are available in the Gene Expression Omnibus (GEO) under accession number GSE210649, while the EX-GF RNA-seq dataset was newly generated in the present study and have been deposited in the GEO under accession number GSE303797. The dataset is currently under private status and will be made publicly available upon publication of the peer-reviewed article.

Raw metagenomic sequencing data are available in the NCBI Sequence Read Archive (SRA) under accession number PRJNA1415844.

Metabolomics data have been deposited in the MassIVE repository (UCSD) under accession number MSV000101376. The dataset is currently under private status and will be made publicly available upon publication of the peer-reviewed article.

The mass spectrometry proteomics data have been deposited to the ProteomeXchange Consortium via the PRIDE (61) partner repository with the dataset identifier PXD077879 and 10.6019/PXD077879. The dataset is currently under private status and will be made publicly available upon publication of the peer-reviewed article.

No custom code or scripts were generated or used in this study. All analyses were performed using standard, publicly available software and established pipelines.

## Author Contributions

G.R., M.C., and K.H.-T. conceived and supervised the study. G.R., M.C., K.H.-T., S.C., and D.P. contributed to experimental design and manuscript drafting. M.B. and S.B. were responsible for breeding and maintaining germ-free, gnotobiotic, conventional, and ex-germ-free mice. M.S. and L.W. performed bioinformatic analyses of the metagenomic data. C.C., F.A., and S.O. conducted RNA-seq and downstream bioinformatic analyses of dorsal root ganglia and sciatic nerve samples. S.K., M.F., and K.H.-T. collected all tissue samples and carried out histomorphometric analysis of the median nerves. G.R., D.P., S.F., and G.G. extracted mRNA from dorsal root ganglia and sciatic nerves and conducted qRT-PCR analyses. M.C., S.C., and D.B. performed all analyses related to muscle samples. D.P., S.F., and F.B. performed quantitative assessments of dorsal root ganglia, measurements of myelin lamellae and nodes of Ranvier length. C.G. and D.C. carried out proteomic analyses on muscle tissues. K.K. performed targeted metabolomic analysis on serum samples. All authors contributed to data interpretation, discussed the results, and reviewed and edited the manuscript.

## Classification

Biological sciences/Neuroscience

## References

1. Y. Chen, J. Zhou, L. Wang, Role and Mechanism of Gut Microbiota in Human Disease. Front Cell Infect Microbiol 11, 625913 (2021).

2. S. Ahlawat, Asha, K. K. Sharma, Gut-organ axis: a microbial outreach and networking. Lett Appl Microbiol 72, 636–668 (2021).

3. P. Mary, L. Servais, R. Vialle, Neuromuscular diseases: Diagnosis and management. Orthop Traumatol Surg Res 104, S89–S95 (2018).

4. K. Hou et al., Microbiota in health and diseases. Signal Transduct Target Ther 7, 135 (2022).

5. J. A. Gilbert et al., Current understanding of the human microbiome. Nat Med 24, 392–400 (2018).

6. J. M. Brown, S. L. Hazen, Targeting of microbe-derived metabolites to improve human health: The next frontier for drug discovery. J Biol Chem 292, 8560–8568 (2017).

7. M. Cescon et al., Gut microbiota depletion delays somatic peripheral nerve development and impairs neuromuscular junction maturation. Gut Microbes 16, 2363015 (2024).

8. S. Lahiri et al., The gut microbiota influences skeletal muscle mass and function in mice. Sci Transl Med 11 (2019).

9. G. Li, B. Jin, Z. Fan, Mechanisms Involved in Gut Microbiota Regulation of Skeletal Muscle. Oxid Med Cell Longev 2022, 2151191 (2022).

10. G. V. Michailov et al., Axonal neuregulin-1 regulates myelin sheath thickness. Science 304, 700–703 (2004).

11. C. Taveggia et al., Neuregulin-1 type III determines the ensheathment fate of axons. Neuron 47, 681–694 (2005).

12. G. Minty et al., aNMJ-morph: a simple macro for rapid analysis of neuromuscular junction morphology. R Soc Open Sci 7, 200128 (2020).

13. S. C. Bokoliya, Y. Dorsett, H. Panier, Y. Zhou, Procedures for Fecal Microbiota Transplantation in Murine Microbiome Studies. Front Cell Infect Microbiol 11, 711055 (2021).

14. R. L. Friede, T. Samorajski, Myelin formation in the sciatic nerve of the rat. A quantitative electron microscopic, histochemical and radioautographic study. J Neuropathol Exp Neurol 27, 546–570 (1968).

15. H. D. Webster, The geometry of peripheral myelin sheaths during their formation and growth in rat sciatic nerves. J Cell Biol 48, 348–367 (1971).

16. B. Garbay, A. M. Heape, F. Sargueil, C. Cassagne, Myelin synthesis in the peripheral nervous system. Prog Neurobiol 61, 267–304 (2000).

17. S. Delgado-Ocana, S. Cuesta, From microbes to mind: germ-free models in neuropsychiatric research. mBio 15, e0207524 (2024).

18. A. Castillo-Ruiz et al., Brain effects of gestating germ-free persist in mouse neonates despite acquisition of a microbiota at birth. Front Neurosci 17, 1130347 (2023).

19. A. E. Hoban et al., Regulation of prefrontal cortex myelination by the microbiota. Transl Psychiatry 6, e774 (2016).

20. S. Belin et al., Neuregulin 1 type III improves peripheral nerve myelination in a mouse model of congenital hypomyelinating neuropathy. Hum Mol Genet 28, 1260–1273 (2019).

21. C. Scapin et al., Enhanced axonal neuregulin-1 type-III signaling ameliorates neurophysiology and hypomyelination in a Charcot-Marie-Tooth type 1B mouse model. Hum Mol Genet 28, 992–1006 (2019).

22. J. Hong et al., PMP2 regulates myelin thickening and ATP production during remyelination. Glia 72, 885–898 (2024).

23. K. Nay et al., Gut bacteria are critical for optimal muscle function: a potential link with glucose homeostasis. Am J Physiol Endocrinol Metab 317, E158–E171 (2019).

24. R. Qi et al., The intestinal microbiota contributes to the growth and physiological state of muscle tissue in piglets. Sci Rep 11, 11237 (2021).

25. Y. Arai, M. Osawa, Y. Fukuyama, Muscle CT scans in preclinical cases of Duchenne and Becker muscular dystrophy. Brain Dev 17, 95–103 (1995).

26. M. Torriani et al., Lower leg muscle involvement in Duchenne muscular dystrophy: an MR imaging and spectroscopy study. Skeletal Radiol 41, 437–445 (2012).

27. M. van Putten et al., Natural disease history of the D2-mdx mouse model for Duchenne muscular dystrophy. FASEB J 33, 8110–8124 (2019).

28. T. R. Valentino et al., Dysbiosis of the gut microbiome impairs mouse skeletal muscle adaptation to exercise. J Physiol 599, 4845–4863 (2021).

29. M. Spinazzi et al., Myotilin gene duplication causing late-onset myotilinopathy. Eur J Neurol 32, e70029 (2025).

30. P. Salmikangas et al., Myotilin, the limb-girdle muscular dystrophy 1A (LGMD1A) protein, cross-links actin filaments and controls sarcomere assembly. Hum Mol Genet 12, 189–203 (2003).

31. K. Wadmore, A. J. Azad, K. Gehmlich, The Role of Z-disc Proteins in Myopathy and Cardiomyopathy. Int J Mol Sci 22 (2021).

32. J. E. Poort, M. B. Rheuben, S. M. Breedlove, C. L. Jordan, Neuromuscular junctions are pathological but not denervated in two mouse models of spinal bulbar muscular atrophy. Hum Mol Genet 25, 3768–3783 (2016).

33. L. Du, R. Qi, J. Wang, Z. Liu, Z. Wu, Indole-3-Propionic Acid, a Functional Metabolite of Clostridium sporogenes, Promotes Muscle Tissue Development and Reduces Muscle Cell Inflammation. Int J Mol Sci 22 (2021).

34. E. Serger et al., The gut metabolite indole-3 propionate promotes nerve regeneration and repair. Nature 607, 585–592 (2022).

35. H. Zhang et al., Indole-3-propionic acid promotes Schwann cell proliferation following peripheral nerve injury by activating the PI3K/AKT pathway. Neurotherapeutics 22, e00578 (2025).

36. J. M. Chen et al., Microbiota-derived IPA mitigates post-stroke neuroinflammation by inhibiting TREM2-dependent pyroptosis. J Neuroinflammation 10.1186/s12974-025-03660-8 (2026).

37. Felasa working group on revision of guidelines for health monitoring of rodents rabbits et al., FELASA recommendations for the health monitoring of mouse, rat, hamster, guinea pig and rabbit colonies in breeding and experimental units. Lab Anim 48, 178–192 (2014).

38. M. Basic et al., Monitoring and contamination incidence of gnotobiotic experiments performed in microisolator cages. Int J Med Microbiol 311, 151482 (2021).

39. P. Ewels, M. Magnusson, S. Lundin, M. Kaller, MultiQC: summarize analysis results for multiple tools and samples in a single report. Bioinformatics 32, 3047–3048 (2016).

40. J. Lu et al., Metagenome analysis using the Kraken software suite. Nat Protoc 17, 2815–2839 (2022).

41. E. Paradis, K. Schliep, ape 5.0: an environment for modern phylogenetics and evolutionary analyses in R. Bioinformatics 35, 526–528 (2019).

42. A. Dobin et al., STAR: ultrafast universal RNA-seq aligner. Bioinformatics 29, 15–21 (2013).

43. Y. Liao, G. K. Smyth, W. Shi, featureCounts: an efficient general purpose program for assigning sequence reads to genomic features. Bioinformatics 30, 923–930 (2014).

44. M. D. Robinson, D. J. McCarthy, G. K. Smyth, edgeR: a Bioconductor package for differential expression analysis of digital gene expression data. Bioinformatics 26, 139–140 (2010).

45. A. Subramanian et al., Gene set enrichment analysis: a knowledge-based approach for interpreting genome-wide expression profiles. Proc Natl Acad Sci U S A 102, 15545–15550 (2005).

46. Z. Gu, R. Eils, M. Schlesner, Complex heatmaps reveal patterns and correlations in multidimensional genomic data. Bioinformatics 32, 2847–2849 (2016).

47. K. Haastert-Talini et al., Chitosan tubes of varying degrees of acetylation for bridging peripheral nerve defects. Biomaterials 34, 9886–9904 (2013).

48. M. Stossel, L. Rehra, K. Haastert-Talini, Reflex-based grasping, skilled forelimb reaching, and electrodiagnostic evaluation for comprehensive analysis of functional recovery-The 7-mm rat median nerve gap repair model revisited. Brain Behav 7, e00813 (2017).

49. M. Moriggi et al., Characterization of Proteome Changes in Aged and Collagen VI-Deficient Human Pericyte Cultures. Int J Mol Sci 25 (2024).

50. J. R. Wisniewski, A. Zougman, N. Nagaraj, M. Mann, Universal sample preparation method for proteome analysis. Nat Methods 6, 359–362 (2009).

51. J. Cox, M. Mann, MaxQuant enables high peptide identification rates, individualized p.p.b.-range mass accuracies and proteome-wide protein quantification. Nat Biotechnol 26, 1367–1372 (2008).

52. J. Cox et al., Accurate proteome-wide label-free quantification by delayed normalization and maximal peptide ratio extraction, termed MaxLFQ. Mol Cell Proteomics 13, 2513–2526 (2014).

53. S. Tyanova et al., The Perseus computational platform for comprehensive analysis of (prote)omics data. Nat Methods 13, 731–740 (2016).

54. A. Kramer, J. Green, J. Pollard, Jr., S. Tugendreich, Causal analysis approaches in Ingenuity Pathway Analysis. Bioinformatics 30, 523–530 (2014).

55. A. Waisman, A. M. Norris, M. Elias Costa, D. Kopinke, Automatic and unbiased segmentation and quantification of myofibers in skeletal muscle. Sci Rep 11, 11793 (2021).

56. S. Reiter et al., Development of a Highly Sensitive Ultra-High-Performance Liquid Chromatography Coupled to Electrospray Ionization Tandem Mass Spectrometry Quantitation Method for Fecal Bile Acids and Application on Crohn’s Disease Studies. J Agric Food Chem 69, 5238–5251 (2021).

57. D. Hacker et al., Exclusive enteral nutrition initiates individual protective microbiome changes to induce remission in pediatric Crohn’s disease. Cell Host Microbe 32, 2019–2034 e2018 (2024).

58. J. Han, K. Lin, C. Sequeira, C. H. Borchers, An isotope-labeled chemical derivatization method for the quantitation of short-chain fatty acids in human feces by liquid chromatography-tandem mass spectrometry. Anal Chim Acta 854, 86–94 (2015).

59. E. Thiele Orberg et al., Bacteria and bacteriophage consortia are associated with protective intestinal metabolites in patients receiving stem cell transplantation. Nat Cancer 5, 187–208 (2024).

60. M. Hemmati et al., Development of a Global Metabo-Lipid-Prote-omics Workflow to Compare Healthy Proximal and Distal Colonic Epithelium in Mice. J Proteome Res 23, 3124–3140 (2024).

61. Y. Perez-Riverol et al., The PRIDE database at 20 years: 2025 update. Nucleic Acids Res 53, D543–D553 (2025).

