## Supplementary Figures for "Post-Weaning Gut Microbiota Colonization Reveals Divergent Recovery of Skeletal Muscle and Peripheral Nerves"

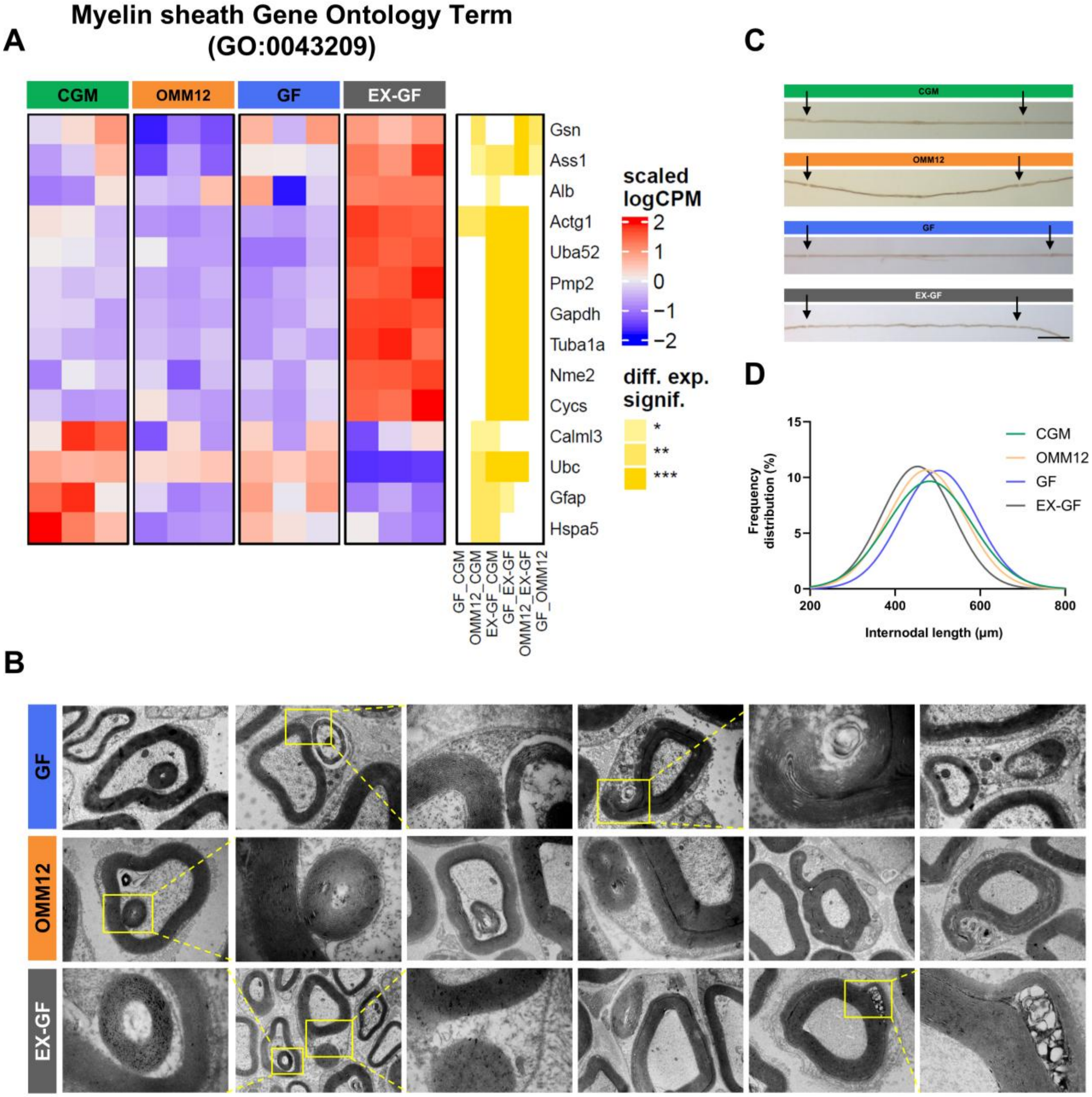


**Supplementary figure S1. Myelin and Schwann cell–axon abnormalities.** (A) Heatmap displaying scaled log-expression values of differentially expressed genes associated with the Gene Ontology (GO) term "myelin sheath" (GO:0043209). Red and blue indicate high and low expression, respectively. Heatmap on the right show the significance value in each pairwise comparisons (*p ≤ 0.05; **p ≤ 0.01; ***p ≤ 0.001). (B) Representative transmission electron microscopy (TEM) images illustrating structural abnormalities in Schwann cell–axon units. (C) Representative light microscopy images of teased ulnar nerve fibers from each group. Black arrows indicate two consecutive nodes of Ranvier. Scale bar: 100 µm. (D) Percentile distribution of internodal length fitted with Gaussian regression curves (equation not shown).

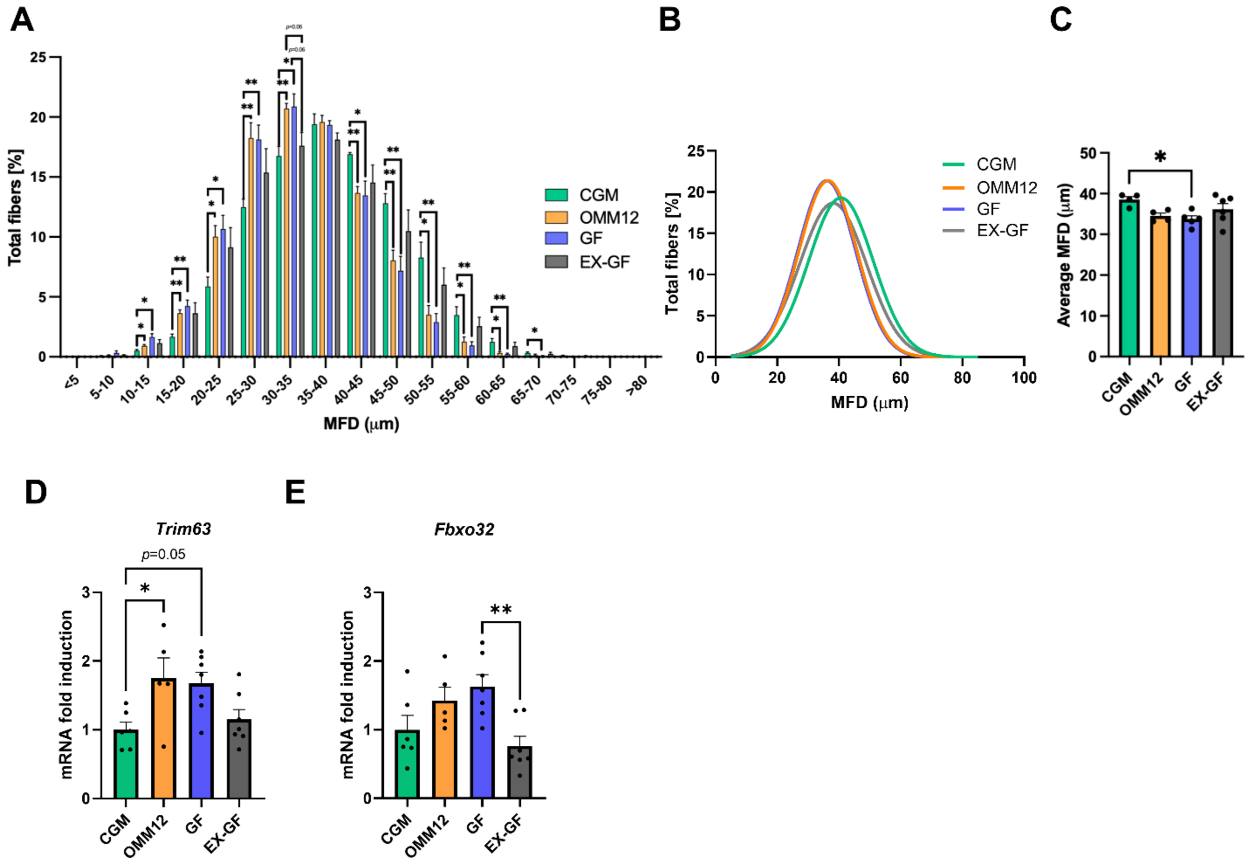


**Supplementary figure S2. Muscle atrophy analysis in GA muscles.** (A) Comparison of MFD distribution among CGM, OMM12, GF and EX-GF GA muscles. (*p ≤ 0.05; **p ≤ 0.01; one-way ANOVA test with Tukey's post hoc test for multiple comparisons; n=3 mice, each group). (B) Graphical representation of MFD distribution among myofibers in GA muscles from CGM, OMM12, GF and EX-GF mice. Curves were fitted to data using non-linear regression (Gaussian). (C) Quantification of average myofiber minimum Feret's diameter in GA muscles from CGM, OMM12, GF and EX-GF mice (*p ≤ 0.05; one-way ANOVA test with Tukey's post hoc test for multiple comparisons; n=3 mice, each group). (D, E) qRT-PCR analysis of transcripts coding for the atrogenes Trim63 (D), Fbxo32 (E) in GA muscle (*p ≤ 0.05; **p ≤ 0.01; one-way ANOVA with post hoc Tukey's multiple comparisons test; CGM, n=8; OMM12, n=5; GF, EX-GF n=7 mice). All data are depicted as mean±SEM.

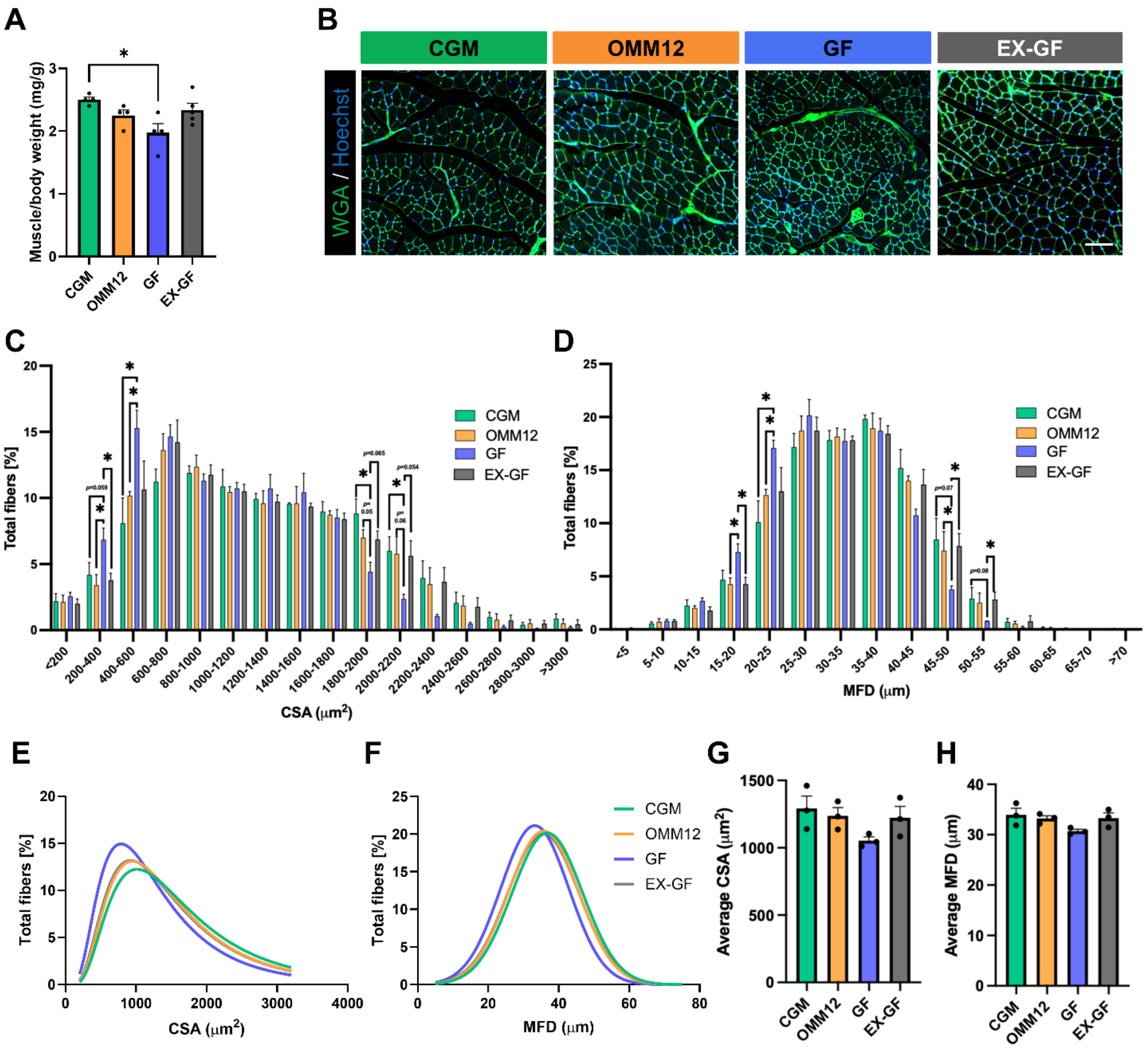


**Supplementary figure S3. Morphometric and molecular analysis of skeletal muscle atrophy in tibialis anterior muscle.** (A) Tibialis anterior (TA) muscle weight normalized by mice body weight from CGM, OMM12, GF and EX-GF mice (*p ≤ 0.51; ordinary one-way ANOVA with post hoc Tukey's multiple comparisons test; CGM, OMM12, GF, n=4; EX-GF, n=5). (B) Representative fluorescence micrographs of TA cross-sections from CGM, OMM12, GF and EX-GF mice, stained with fluorophore-conjugated wheat germ agglutinin (WGA, green) and Hoechst (blue). Scale bar: 100 µm. (C, D) Quantification of average myofiber cross- sectional area (CSA) (C) and minimum Feret's diameter (MFD) (D) in TA muscles from CGM, OMM12, GF and EX-GF mice (one-way ANOVA test with Tukey's post hoc test for multiple comparisons; n=3 mice, each group). (E, F) Comparison of myofiber CSA (E) and MFD (F) distribution among CGM, OMM12, GF and EX-GF TA muscles. (*p ≤ 0.05; multiple unpaired two-tailed Student's t-tests; n=3 mice, each group). (G, H) Graphical representation of CSA (G) and MFD (H) distribution among myofibers in TA muscles from CGM, OMM12, GF and EX-GF mice. Curves were fitted to data using nonlinear regression (Lognormal). All data are represented as mean±SEM.

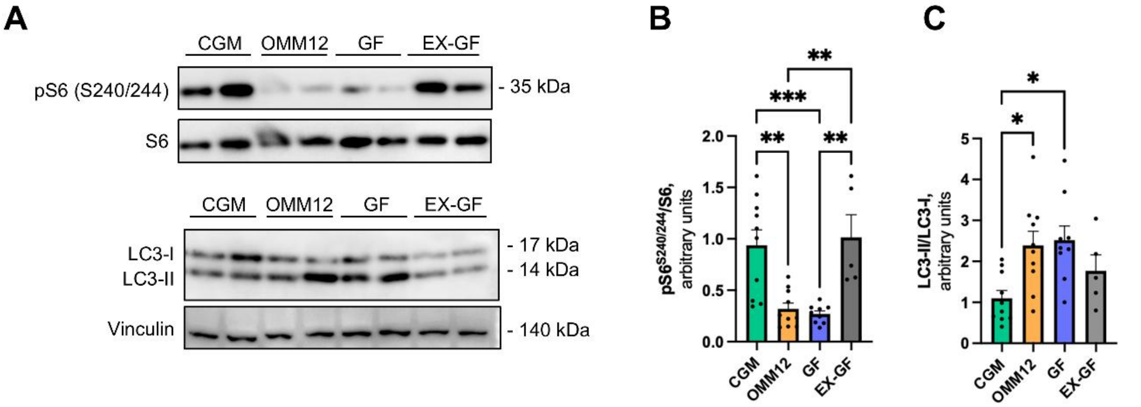


**Supplementary figure S4. Western blot analysis of S6 phosphorylation and LC3 lipidation in TA muscles.** (A) Representative images of western blot analysis of LC3 lipidation and S6 phosphorylation in total protein lysates of TA muscles from CGM, OMM12, GF and EX-GF mice. (B, C) Densitometric quantifications of pS6 normalized to total S6 (A) and of LC3-II normalized to LC3-I (B) (*p ≤ 0.05; **p ≤ 0.01; ***p ≤ 0.001; one-way ANOVA with post hoc Tukey's multiple comparisons test; CGM, OMM12, n=10; GF, n=9; EX-GF, n=5 mice). All data are depicted as mean±SEM.

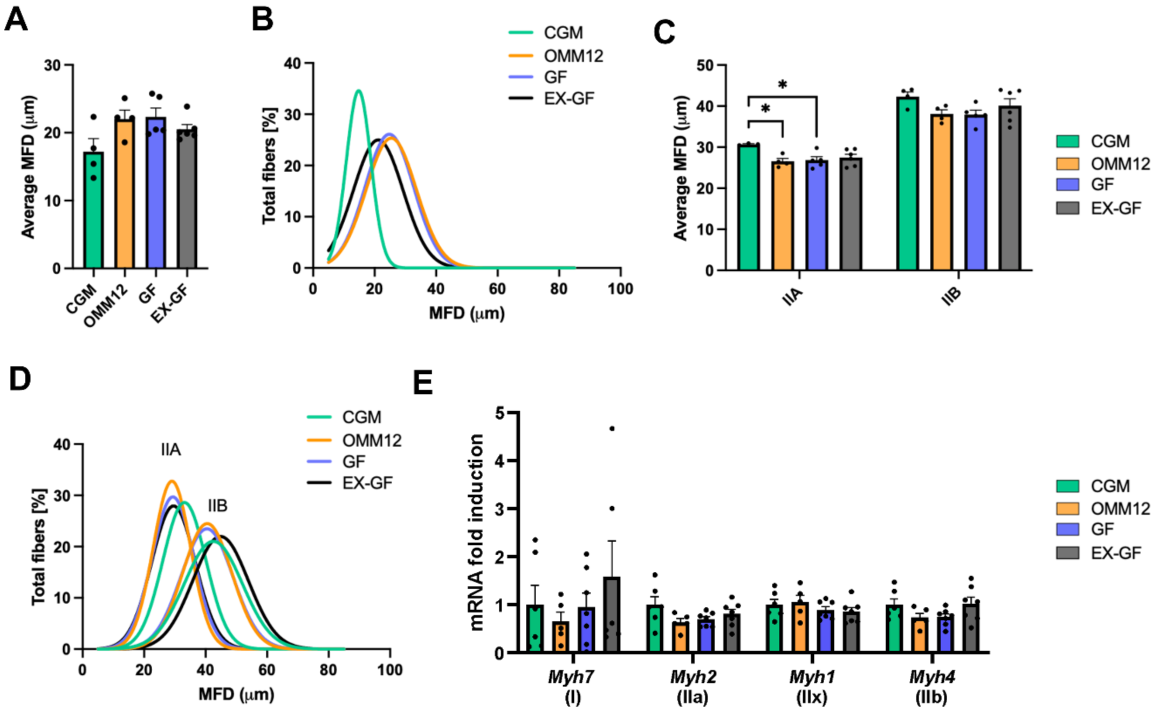


**Supplementary figure S5. Analysis of MFD in GA muscles according to fiber type.** (A) Quantification of the average type I fiber MFD in GA muscles from CGM, OMM12, GF and EX-GF mice (one-way ANOVA test with Tukey's post hoc test for multiple comparisons; CGM, OMM12, n=4; GF n=5; EX-GF n=6). (B) Graphical representation of type I fiber MFD distribution in GA muscles from the experimental groups. Curves were fitted to data using non-linear regression (Gaussian). (C) Quantification of the average type IIA and IIB fibers MFD in GA muscles from CGM, OMM12, GF and EX-GF mice (*$p \leq 0.05$; one-way ANOVA test with Tukey's post hoc test for multiple comparisons; CGM, OMM12 n=4; GF n=5; EX-GF n=6). (D) Graphical representation of type IIA and IIB fibers MFD distribution in GA muscles from the experimental groups. Curves were fitted to data using non-linear regression (Gaussian). (E) qRT-PCR quantification of the levels of transcripts for different myosin heavy chain (MHC) isoforms in GA muscles of CGM, OMM12, GF and EX-GF mice (ordinary one-way ANOVA with post hoc Tukey's multiple comparisons test or Kruskal-Wallis with post hoc Dunn's test for multiple comparisons; CGM n=8; OMM12 n=5; GF n=7; EX-GF n=7). All data are shown as mean±SEM.

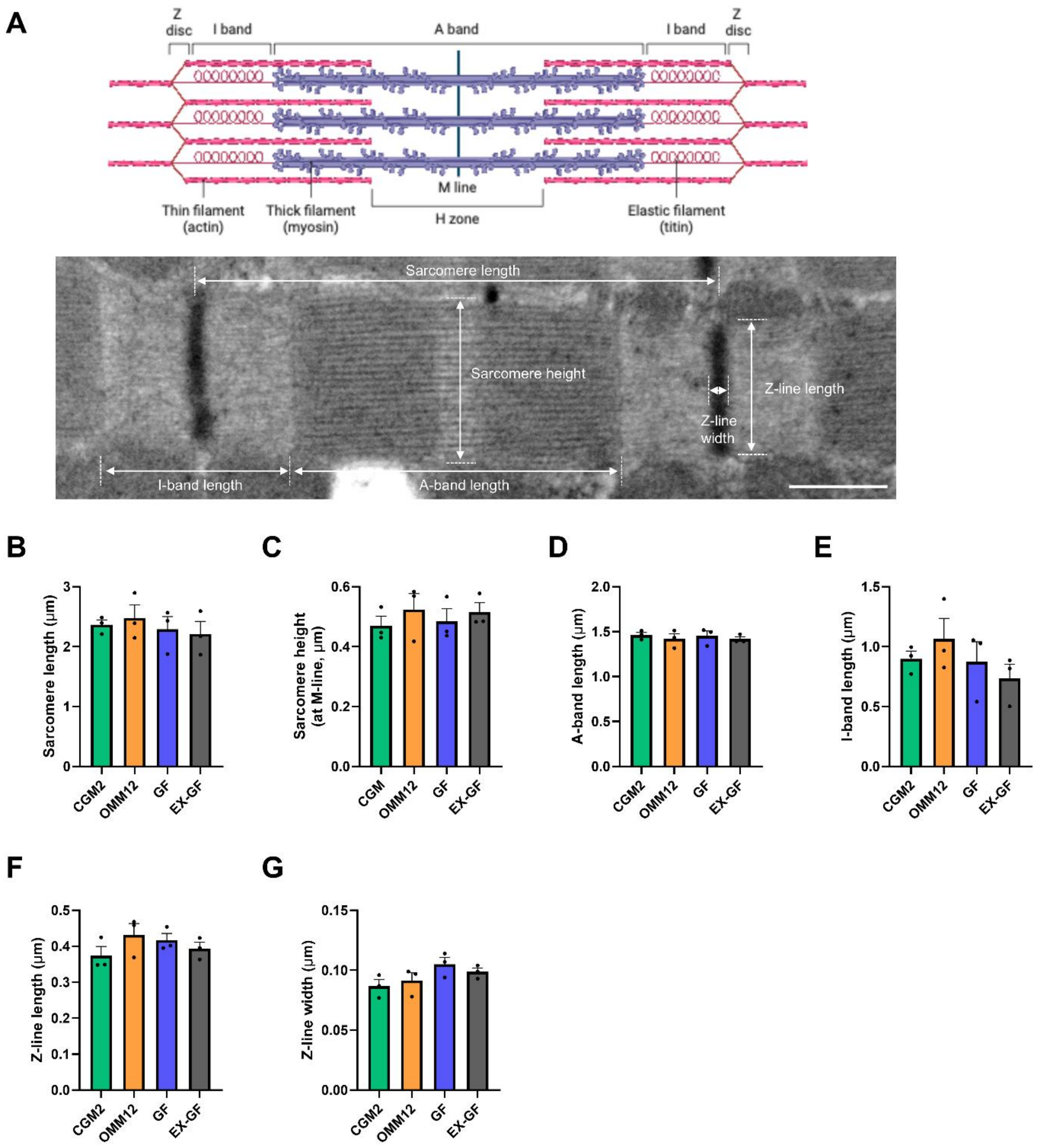


**Supplementary figure S6: Evaluation of sarcomeric morphometry in diaphragm muscles.** (A) Upper panel: schematic illustration of the sarcomere structure showing the architectural arrangement of its main constituents (created with BioRender.com). Bottom panel: representative transmission electron microscopy image of generic sarcomere ultrastructure, with indication of the parameters measured in the following morphometric analysis. Scale bar: 500 nm. (B-G) Quantitative analysis of mean resting sarcomere length (B) and height (C), A-band (D) and I-band length (E), Z-band length (F) and width (G) measured from TEM images, as in (A), of longitudinal sections of diaphragms from CGM, OMM12, GF, and EX-GF mice (one-way ANOVA test with Tukey's post hoc test for multiple comparisons; n=3 each group). All data are depicted as mean±SEM.

**A**

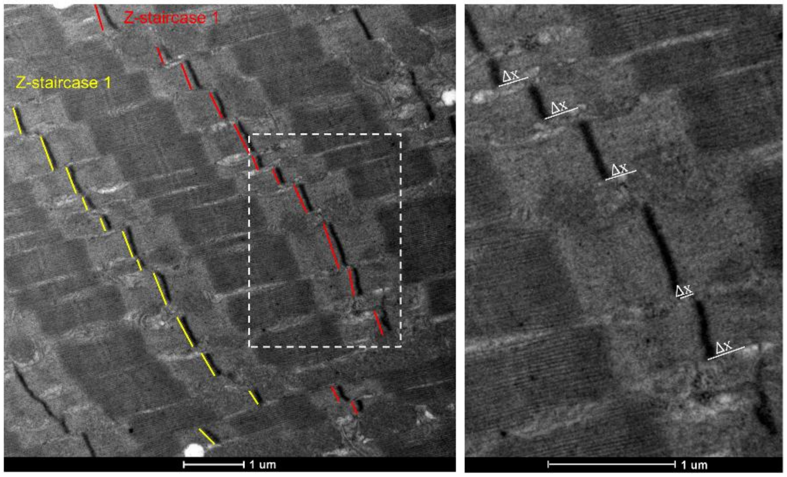


**B**

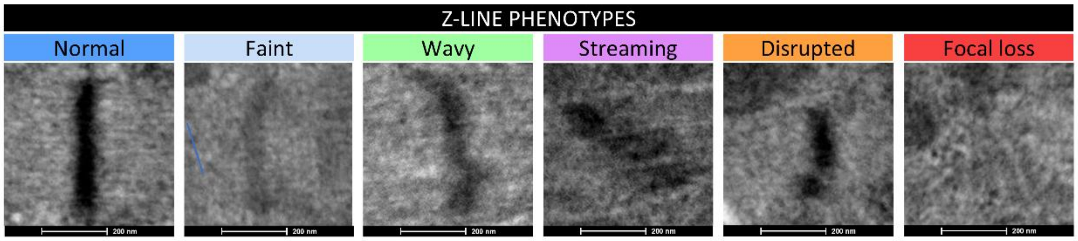


**Supplementary figure S7. Analysis of Z-line misalignment and structural phenotype.** (A) Schematic representation of Z-line misalignment measurement in TEM images of longitudinal sections of diaphragm muscles. The dotted white rectangle indicates the area that is shown at higher magnification in the right panel. Z-line displacement (Δx), defined as the lateral (x-axis) distance between Z-lines of adjacent myofibrils, was measured along continuous Z-line rows ("Z-staircases") and then averaged across multiple Z-staircases. Scale bar: 1 µm. (B) Representative TEM images of different Z-line phenotypes classified based on Z-line ultrastructural appearance in longitudinal diaphragm sections: normal (continuous, sharp and dark Z-lines), faint (Z-lines with reduced electron density and poor contrast), wavy (undulated and irregular Z-lines), streaming (blurred and widened Z-line), disrupted (fragmented and discontinuous Z-lines), focal loss (complete absence of Z-lines in a localized region). Scale bar: 200 nm.

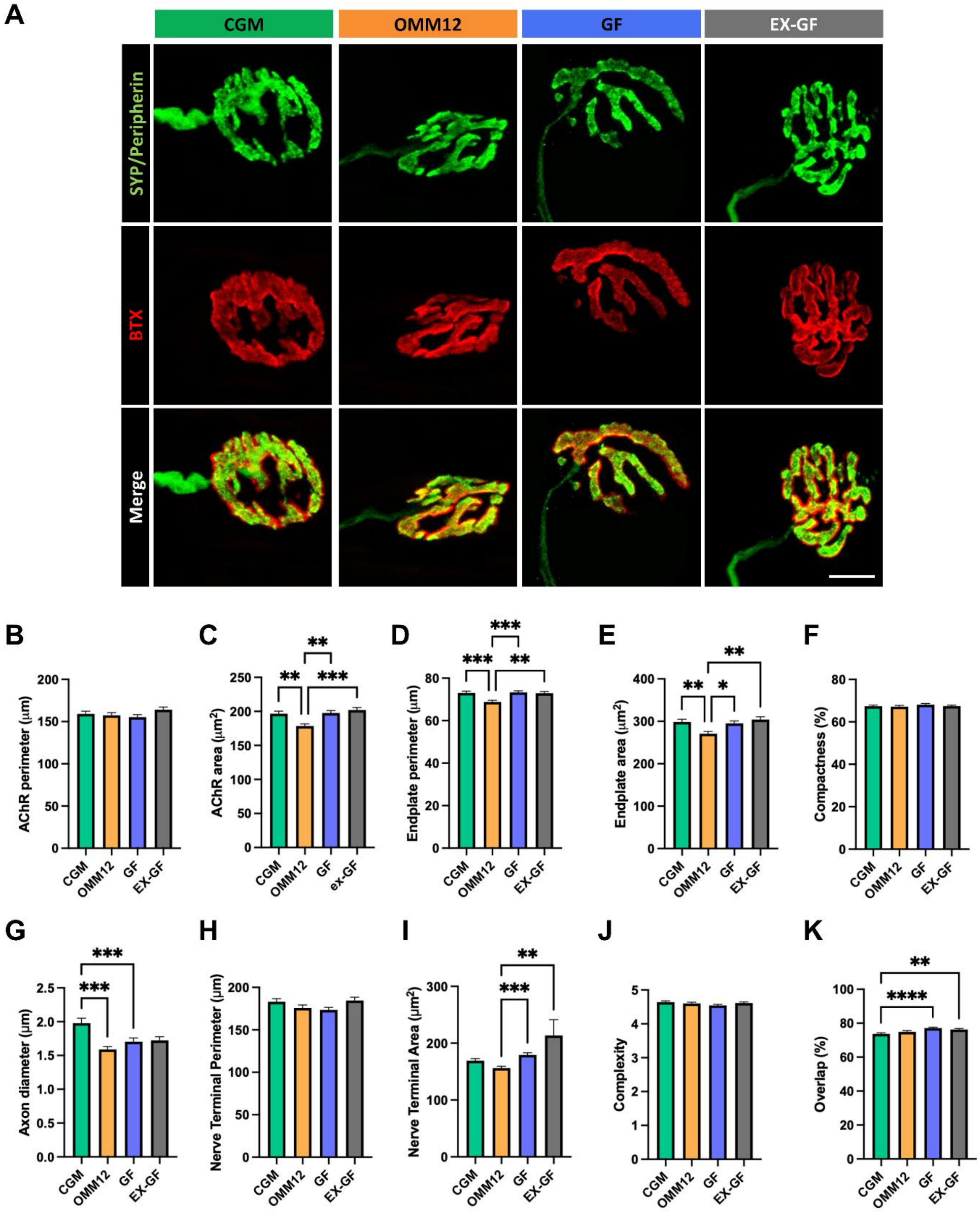


**Supplementary figure S8. Morphometric analysis of NMJ pre- and post-synaptic terminals in diaphragm muscles.** (A) Representative confocal images of NMJs labeled with α-bungarotoxin (BTX, red) and antibodies to synaptophysin (SYP) and peripherin (green) in diaphragm muscles from CGM, OMM12, GF, and EX-GF mice. Scale bar: 10 μm. (B-F) Quantitative analysis of postsynaptic parameters, such as acetylcholine receptor (AChR) perimeter (B), AChR area (C), endplate perimeter (D), endplate area (E) and compactness (F) of NMJs from CGM,

OMM12, GF, and EX-GF mice (*p ≤ 0.05; **p ≤ 0.01; ***p ≤ 0.001; Kruskal-Wallis with Dunn's post hoc test for multiple comparisons; CGM, n=270; OMM12 and EX-GF, n=279; GF, n=304 NMJs from 4 mice, each group). (G) Quantification of axon diameter in NMJs from CGM, OMM12, GF, and EX-GF mice (***p ≤ 0.001; Kruskal-Wallis with Dunn's post hoc test for multiple comparisons; CGM n=128; OMM12 n=134; GF n=179; EX-GF n=103 NMJs from 4 mice, each group). (H-K) Quantitative analysis of presynaptic parameters, such as nerve terminal perimeter (H), nerve terminal area (I), and complexity (J) in NMJs from CGM, OMM12, GF, and EX-GF mice (**p ≤ 0.01; ***p ≤ 0.001; Kruskal-Wallis with Dunn's post hoc test for multiple comparisons; CGM n=241; OMM12 n=246; GF n=300; EX-GF n=262 NMJs from 4 mice, each group). (K) Measure of the degree of overlap between the presynaptic and the postsynaptic staining in NMJs from CGM, OMM12, GF, and EX-GF mice (**p ≤ 0.01; ****p ≤ 0.0001; Kruskal-Wallis with Dunn's post hoc test for multiple comparisons; CGM n=241; OMM12 n=246; GF n=300; EX-GF n=262 NMJs from 4 mice, each group). All data are expressed as mean±SEM.
